# iASPP contributes to cortex rigidity, astral microtubule capture and mitotic spindle positioning

**DOI:** 10.1101/870998

**Authors:** Aurélie Mangon, Danièle Salaün, Mohamed Lala Bouali, Sabine Quitard, Daniel Isnardon, Stéphane Audebert, Pierre-Henri Puech, Pascal Verdier-Pinard, Ali Badache

## Abstract

The microtubule plus-end binding protein EB1 is the core of a complex protein network which regulates microtubule dynamics during important biological processes such as cell motility and mitosis. We found that iASPP, an inhibitor of p53 and predicted regulatory subunit of the PP1 phosphatase, associates with EB1 at microtubule plus-ends via a SxIP motif. iASPP silencing or mutation of the SxIP motif led to defective microtubule capture at the leading edge of migrating cells, and at the cortex of mitotic cells leading to abnormal positioning of the mitotic spindle. These effects were recapitulated by the knockdown of Myosin-Ic (Myo1c), identified as a novel partner of iASPP. Moreover, iASPP or Myo1c knockdown cells failed to round up during mitosis because of defective cortical rigidity. We propose that iASPP, together with EB1 and Myo1c, contributes to mitotic cell cortex rigidity, allowing astral microtubule capture and appropriate positioning of the mitotic spindle.

## Introduction

Regulation of microtubule dynamics is central to major biological processes, including cell division and motility, that occur during normal development, but also upon tumor formation and dissemination.

Microtubules are nucleated at microtubule organizing centers, often the centrosomes, and elongate through sequential phases of growth and shortening, until reaching structures, such as the cell cortex or kinetochores, where they will be captured and stabilized. Microtubule stabilization at the cell cortex is required for oriented trafficking of intracellular vesicles and directed cell motility^1–3^. Astral microtubule capture during mitosis is required for mitotic spindle orientation and positioning of the mitotic spindle. In monolayered epithelia, orientation of the mitotic spindle parallel to the basal surface allows both daughter cells to remain in the plane of the monolayer, whereas orthogonal division is required for daughter cell differentiation and epithelia stratification. In symmetric cell division, placement of the mitotic spindle at the cell center and orientation along the future axis of cell division is required for equal distribution of the cellular material to the daughter cells^4^. During asymmetric division, positioning of the spindle away from the cell center allows divergent cell fate of the daughter cells^5,6^. Spindle orientation and positioning are driven by pulling forces, exerted on astral microtubules emanating from spindle poles by the cortically-anchored minus-end directed motor complex dynein-dynactin8,9. Anchoring of the motor proteins involves a well conserved ternary complex, including α-subunits of heterotrimeric G proteins, the G protein regulator LGN (Leu-Gly-Asp repeat protein), and the nuclear and mitotic apparatus protein (NuMA), which has been shown to interact with the dynein-dynactin complex and microtubules^7,8^. Other proteins associated with the actomyosin cortex and/or the plasma membrane, including Dlg (discs large), afadin and MISP (mitotic interactor and substrate of Plk1) were also found to regulate the positioning of the spindle in metaphase^9^, most of which were proposed to function via the control of LGN and NuMA localization. The ERM (ezrin/radixin/moesin) proteins are actin-membrane linkers that were shown to regulate spindle orientation and positioning, but it was not clear if this involved the control of LGN-NuMA localization^10^ or cortex properties^11,12^. Of note, the mechanisms of spindle centering and orientation are rarely distinguished in these studies and it is generally assumed that the two processes are controlled by the same cortical complexes.

In cells, microtubule function is modulated by microtubule-associated proteins, including plus-end tracking proteins (+TIPs), a large and diverse family of proteins that share the ability to bind growing microtubules plus-ends^13^. EB1 is a hub in the complex network of +TIPs. It directly interacts with microtubule plus-ends and recruits many other proteins harboring a SxIP or CAP-Gly motif, to control microtubule dynamics and mediate their association with the cell cortex. Systematic investigations of EB1 network of protein-protein interactions, implemented by several groups including ours, have revealed numerous potential EB1 partners^14,15^. Deeper investigation of these protein networks is required to delineate macromolecular functional units that regulate specific aspects of microtubule functions^14^.

Here, we investigated an unexplored connection between EB1 and the protein iASPP (inhibitor of ASPP) and its impact on microtubule-dependent biological processes. iASPP is a member of the ASPP (apoptosis-stimulating protein of p53) family of proteins^16^, which share the ability to interact with the p53 tumor suppressor via their C-terminal SH3 domain, to modulate the expression of pro-apoptotic target genes. ASPP proteins also interacts with PP1 and, as such, might act as PP1 targeting subunits^17^. While earlier studies focused on the ability of iASPP to inhibit p53^16,18^, more recent work has identified new functions of iASPP, associated with the Pro-rich region. For instance, iASPP interacts with desmine and desmoplakin in cardiomyocytes to sustain desmosome integrity^19^ and with the Nrf2 transcription factor regulator Keap1, to promote antioxidative signaling^20^. We report here that iASPP interacts with EB1 via a typical SxIP motif located within the intrinsically disordered region. Remarkably, disturbing iASPP-EB1 interaction led to defective mitotic spindle positioning, as a consequence of asymmetric anchoring of astral microtubules. This is also recapitulated by inhibiting the expression of the motor protein Myosin-Ic (Myo1c), that we identified as a partner of iASPP. Finally, we show that both iASPP and Myo1c promote cell cortex stiffness. Thus, we propose that iASPP acts together with EB1 and Myo1c, but independently of p53 or PP1, to control cortical rigidity, microtubule capture and mitotic spindle positioning.

## Results

### iASPP engages in multiple intramolecular and intermolecular interactions

Regulation of microtubule function is critical for major biological processes including cell motility and cell division. In order to better understand how microtubule function is controlled, we have implemented a systematic investigation of the protein interaction network of EB1, in SKBr3 cells stably expressing EB1-GFP^14^. We identified iASPP as a new partner of EB1. We confirmed that iASPP co-immunoprecipitated with EB1 in various cell lines (Fig. 1a). Using a proximity biotinylation assay^21^, we verified that the iASPP-EB1 interaction also occurred *in situ*, before cells were lysed (Fig. 1b). Finally, we showed that GFP-tagged iASPP co-localized with EB1, at microtubules plus-ends (Fig. 1c).

**Figure 1.**
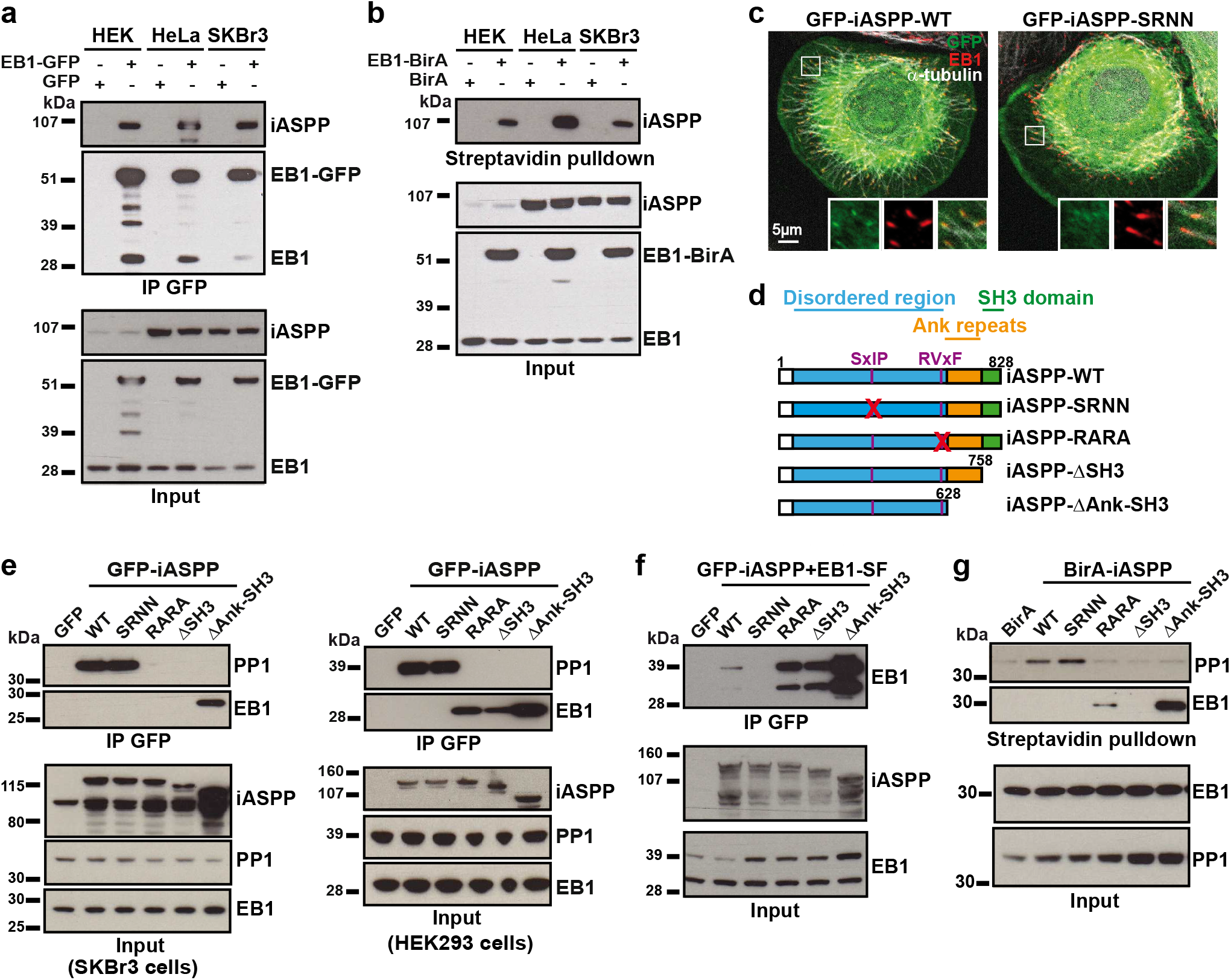
iASPP is a partner of EB1. (**a**) iASPP co-immunoprecipitates with EB1. EB1-GFP or GFP was expressed in the indicated cell lines, before immunoprecipitation with GFP-Trap and Western blotting. (**b**) iASPP is a neighbor of EB1 *in situ*. EB1-BirA was expressed in the indicated cell lines, before addition of biotin, isolation of EB1 proximal (biotinylated) proteins with streptavidin and Western blotting. (**c**) iASPP co-localizes with EB1 at microtubule plus-ends. Microtubules and EB1 were visualized by immunofluorescence in SKBr3 cells expressing GFP-iASPP or GFP-iASPP-SRNN. Inserts: zoomed images (2.5×) of the boxed regions. (**d-g**) Identification of motifs and domains required for EB1 and PP1 binding. iASPP constructs used in the present study are described in (**d**). GFP-tagged (**e,f**), BirA-tagged iASPP (**g**), and Strep-FLAG (SF)-tagged EB1 (**f**) constructs were expressed in SKBr3 (**e,g**) or HEK293F (**e,f**) cells, and co-immunoprecipitation (**e,f**) or biotinylation (**g**) of PP1 and EB1 analyzed by Western blotting.

In order to investigate how iASPP interacts with EB1, but also other partners, we introduced specific modifications in iASPP via directed mutagenesis (Fig. 1d): mutation of a putative EB1-binding SxIP motif (^350^SRIP^353^ to SRNN); mutation of the putative PP1-binding motif^17,22^ (^622^RARL^625^ to RARA); deletion of the C-terminal SH3 domain (iASPP-ΔSH3) required for the interaction with p53^16^; or deletion of the four ankyrin repeats and the SH3 domain (iASPP-ΔAnk-SH3). We then evaluated the ability of the various constructs to interact with EB1 and PP1 by immunoprecipitation, in SKBr3 and HEK293F cells (Fig. 1e). We found that wild type (WT) iASPP interacted strongly with PP1. The interaction was lost when the SH3 domain was absent or the RARL motif mutated. Conversely, the C-terminal region (Cter, region 600-828) of iASPP which included the RARL and SILK motifs^23^, the Ank repeats and the SH3 domain was sufficient for robust association of PP1 with iASPP (Supplementary Fig. 1a). Deletion of the SILK/RARL motifs (Ank-SH3, region 628-828) strongly decreased PP1 binding, whereas the Ank repeats or the SH3 domain alone showed no affinity for PP1 (Supplementary Fig. 1a), confirming that PP1 binds iASPP via multiple cooperative C-terminally located regions.

Upon iASPP pulldown, the interaction with EB1 was not detectable (Fig. 1f), unless we ectopically expressed tagged EB1 (Fig. 1f), indicative of a weaker affinity than PP1. Interaction of iASPP with EB1 (Fig. 1f) and co-localization with EB1 (Fig. 1c) was prevented upon mutation of the SRIP motif. Interestingly, EB1 bound stronger to iASPP constructs that did not interact with PP1 (iASPP-RARA and iASPP-ΔSH3) and even better with the iASPP-ΔAnk-SH3 construct which lacks a large C-terminal fragment (Fig. 1e), indicating that PP1 and the C-terminal region hinder EB1 interaction with iASPP. It is noteworthy that, with the exception of GFP-iASPP-SRNN, all GFP-iASPP constructs co-localized with EB1 comets at microtubules plus-ends (Supplementary Fig. 2). Through proximity biotinylation assays, we confirmed that the patterns of interaction of iASPP constructs with PP1 and EB1, described above, reflected the *in vivo* situation (Fig. 1g).

Of note, we have not been able to detect a significant interaction of iASPP with p53 in the cellular models used in the present study, in accordance with the demonstrated lack of iASPP nuclear localization aside from melanoma cell lines^24^.

A previous study suggested that iASPP constitutes homodimers in the cytoplasm, the N-terminus of a molecule binding the C-terminus of another molecule; and that phosphorylation by cyclin B1/CDK1 (e.g. upon nocodazole treatment) inhibits dimerization allowing entry into the nucleus^24^.

We thus investigated whether N-terminal iASPP interacted with C-terminal iASPP. We expressed a short N-terminal fragment of iASPP (1-290, similar to the one used in Lu et al^24^) or a long N-terminal fragments corresponding to the whole disordered regions (1-628, or 1-599 to prevent overlap with the C-terminal fragment; Fig. 2c); together with a C-terminal fragment: 600-828 which includes the RARL motif and thus strongly binds PP1 or 629-828 that poorly binds PP1. N-terminal fragments and C-terminal fragments were fused to SF and GFP, respectively, for GFP-trap pulldowns of the C-terminal fragments in a first set of experiments (Fig. 2a); and the tags were reversed for pulldowns of the N-terminal fragments in a second set of experiments (Fig. 2d).

**Figure 2.**
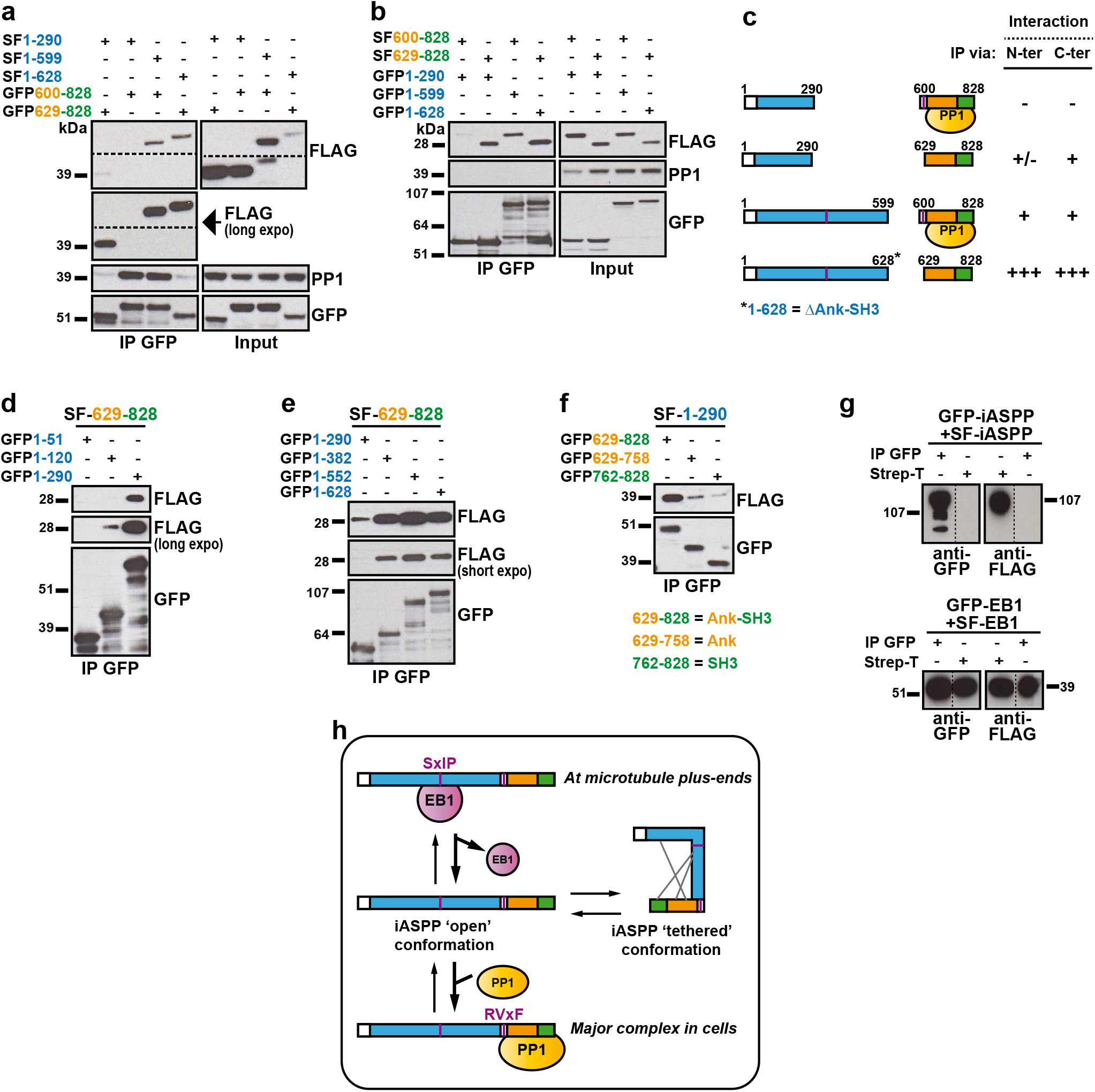
iASPP N-terminal region interacts with iASPP C-terminal region; but iASPP does not form homodimers. (**a-f**) Strep-FLAG (SF)- and GFP-tagged constructs corresponding to the indicated region of iASPP were co-expressed in HEK293F cells, before immunoprecipitation with GFP-Trap and Western blotting of the iASPP fragments and PP1. (**a**) Immunoprecipitation was performed via iASPP C-terminal fragments. (**b**) Immunoprecipitation was performed via iASPP N-terminal fragments. (**c**) Summary of the immunoprecipitation experiments described in (**a**) and (**b**). iASPP disordered region interacts with iASPP C-terminal region; PP1 interferes with the interaction. (**d,e**) Characterization of the N-terminal zones required for interaction with the C-terminal region. (**f**) Characterization of the C-terminal domains required for interaction with the N-terminal region. (**g**) iASPP does not form homodimers. GFP- and SF-tagged iASPP constructs (top) or GFP-tagged and SF-tagged EB1 constructs (bottom) were co-expressed in HEK293F cells, before immunoprecipitation with GFP-Trap or Strep-Tactin beads (Strep-T) and Western blotting with anti-GFP and anti-FLAG antibodies. In contrast to the dimeric protein EB1, GFP-iASPP did not co-precipitate with SF-iASPP. (**h**) Model of iASPP complex organization based on results described in Fig. 1 and 2.

We observed that C-terminal iASPP (region 629-828) interacted modestly with the short N-terminal fragment (1-290) and much better with the longer fragments (1-599/1-628) (Fig. 2a-c). Further molecular mapping indicated that an area located between amino-acids 120 and 290 (Fig. 2d) and another area between amino-acids 290 and 382 (Fig. 2e) had the strongest contributions. On the C-terminal side, the combined contribution of the ankyrin repeats and the SH3 domain was required (Fig. 2f).

Interestingly, when the C-terminal fragment was bound to PP1, its affinity for the N-terminal region was diminished (for constructs 1-599 or 1-628) or even prevented (for construct 1-290) (Fig. 2a-c). Actually, when the C-terminal region was used as bait, we could pull down N-terminal iASPP and PP1 (Fig. 2a). However, when the N-terminal region served as bait (Fig. 2c), we could efficiently pull down the C-terminal region, but not PP1, showing that C-terminal iASPP associated either with N-terminal iASPP or PP1, but not with both simultaneously.

Of note, in our hands, nocodazole treatment did not prevent the interaction between C-terminal iASPP and N-terminal iASPP (Supplementary Fig. 1b). Nocodazole treatment, however, weakened the interaction of iASPP with EB1 (Supplementary Fig. 1c).

To verify if iASPP form homodimers, we co-expressed GFP-tagged and Strep-Flag (SF)-tagged iASPP in HEK293F cells and analyzed the occurrence of co-precipitation. Surprisingly, we detected no associated SF-iASPP into GFP-iASPP immunoprecipitate, and reciprocally, SF-iASPP pulldown revealed no bound GFP-iASPP (Fig. 2g). In a similar experiment using tagged forms of EB1, a protein well-known to form homodimers^25^, EB1-GFP robustly co-precipitated with EB1-SF (Fig. 2g).

Thus, iASPP N-terminal and C-terminal interaction is possible, but does not occur in the context of a homodimer. Altogether, our results suggest a fundamentally different model of iASPP supramolecular organization. iASPP is mostly organized as a complex with PP1. In specific instances, e.g. at microtubule plus-ends, iASPP associates with EB1. We propose that iASPP switches between several conformations: open states in association with PP1 or EB1; and possibly a tethered conformation, thanks to interactions between the folded C-terminal region and the disordered region, the prevalence of which remains to be evaluated (Fig. 2h).

### iASPP contributes to mitotic spindle positioning via its association with EB1

Mitotic microtubules organize a symmetric spindle between centrosomes, which serves to capture and align chromosomes at the cell center. Astral microtubules transmit pulling and pushing forces, generated by evolutionary conserved cortical complexes, to the spindle. As we observed that iASPP interacts with EB1, we evaluated the contribution of iASPP to mitotic microtubule functions. HeLa cells spontaneously align their mitotic spindle parallel to the substrate^9^. We thus used HeLa cells as a model to examine the contribution of iASPP to the organization, positioning and orientation of the mitotic spindle. RNAi-mediated silencing of iASPP (Supplementary Fig. 3) did not significantly affect bipolar spindle formation or the alignment of chromosomes at the metaphase plate (Supplementary Fig. 4a). However, the mitotic spindle often appeared shifted to one side of the cell, instead of being positioned at the cell center (Fig. 3a). We evaluated the distance from the pole to the cortex, on each side of the spindle in iASPP knockdown cells relative to control cells. While the shorter pole-to-cortex distance (d1, Fig. 3b, left) was virtually unchanged, the pole-to-cortex distance on the other side (d2) was significantly increased (Supplementary Fig. 4b). To overcome variability in pole-to-cortex distances due to disparate cell sizes, we calculated the mitotic spindle shift (d2-d1), which should tend towards zero for perfectly centered spindles. In control cells, d1 and d2 distribution was largely overlapping (Supplementary Fig. 4c), and spindle shift around 1 μm (Fig. 3b). iASPP silencing induced a robust shift (close to 3 μm) of the mitotic spindle (Fig. 3b, top right and Supplementary Fig. 4c). As a change in the size of the hemi-spindles might be a confounding factor, we also measured the cortex-to-chromosome plate distance shift (D2-D1) and observed a similar increase of spindle shift upon iASPP knockdown (Fig. 3b, bottom). Comparison of the average control cell and iASPP knockdown cell indicated that the observed spindle shift was the combined consequence of increased cell size and decreased spindle size (Supplementary Fig. 4b, d & e). Thus, our data indicates that iASPP contributes to appropriate mitotic spindle positioning.

**Figure 3.**
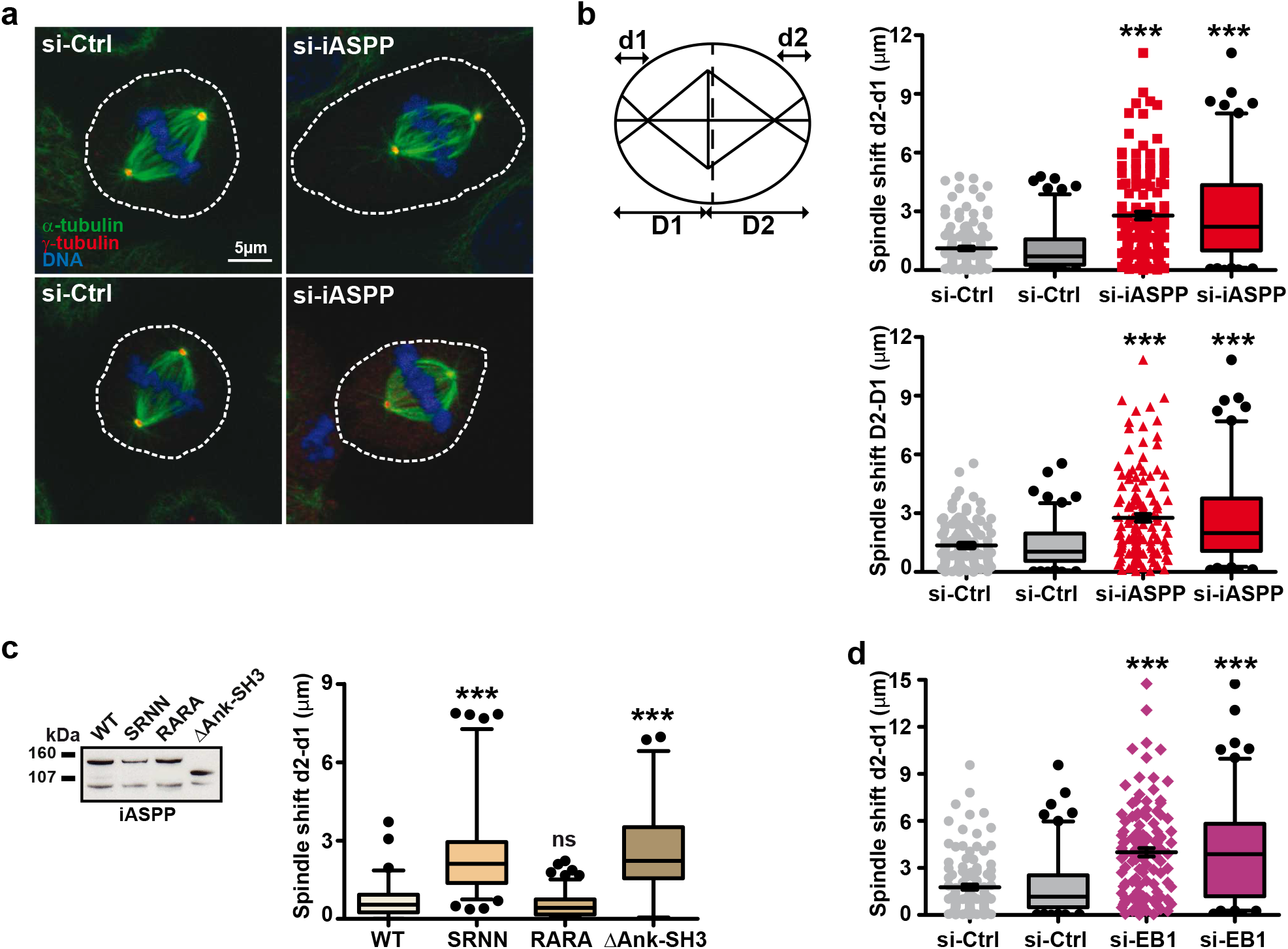
iASPP is required for mitotic spindle positioning in metaphase, via its interaction with EB1. (**a-b**) iASPP silencing induces abnormal mitotic spindle positioning. HeLa cells were transduced with control (Ctrl) or iASPP targeting siRNA. (**a**) Mitotic spindle, centrosomes and chromosomes of metaphasic HeLa cells were visualized by immunofluorescence against α-tubulin and γ-tubulin and DAPI. (**b**) Mitotic spindle positioning was quantified by comparing the distance between cortex and pole (d1 and d2) or cortex and metaphase plate (D1 and D2), on each side of the spindle. (**c**) iASPP defective for EB1 binding induces abnormal spindle positioning. iASPP constructs were stably expressed in HeLa cells. Expression levels were evaluated by Western blotting (*right*) and spindle positioning quantified as above (*left*). (**d**) EB1 silencing induces abnormal spindle positioning. HeLa cells were transduced with Control (Ctrl) or EB1 targeting siRNA and spindle positioning quantified as above. Data were pooled from three independent experiments for a total of 128 and 133 cells for si-Ctrl and si-iASPP respectively (**b**); 94, 90, 72 and 102 cells for WT, SRNN, RARA and ΔAnk-SH3 respectively (**c**) and 120 cells for si-Ctrl and si-EB1 (**d**). Data are represented as scatter plots with mean and SEM; and as box-and-whiskers plots with boxes showing 25^th^ percentile, median and 75^th^ percentile and whiskers indicating the 5^th^ and 95th percentile. *** p<0.001; ns: not significant, relative to si-Ctrl in (**b, d**) or to WT in (**c**), using unpaired t-test with Welch’s correction.

iASPP binds PP1 and EB1. In order to investigate the respective contribution of iASPP-EB1 and iASPP-PP1 interactions to spindle positioning, we derived HeLa cell lines that stably expressed GFP-tagged iASPP constructs (Fig. 3c, left) mutated in motifs and domains important for protein interaction: iASPP-SRNN, iASPP-RARA or iASPP-ΔAnk-SH3, or iASPP-WT as control. All iASPP constructs, except iASPP-SRNN, co-localized with EB1 comets (Supplementary Fig. 5). The analysis of mitotic spindles in metaphase cells revealed that expression of iASPP-SRNN, defective for EB1 binding, induced a large shift in mitotic spindle positioning (Fig. 3c, right). Importantly, cells expressing iASPP-RARA, defective for PP1 binding, displayed well-positioned spindles, clearly showing that PP1 binding is not required for iASPP-dependent spindle positioning. Intriguingly, iASPPΔAnk-SH3 that showed defective PP1 binding, but strongly increased EB1 binding, was also defective for mitotic spindle centering, indicating that the Ank-SH3 domain could be required for this function or that excessive iASPP-EB1 interaction might be detrimental. Comparison of average cells showed a very good overlap between iASPP and iASPP-RARA expressing cells on the one hand, and iASPP-SRNN and iASPPΔAnk-SH3 cells on the other hand (Supplementary Fig. 6), with only the latest showing larger cell size. These results demonstrated that iASPP-EB1 interaction was critical for mitotic spindle positioning. If this is true, one would expect that EB1 inactivation would similarly affect spindle positioning. This was actually the case as EB1 silencing induced a strong shift of the mitotic spindle (Fig. 3d).

### Myo1c is a partner of iASPP involved in spindle positioning

In order to understand the mechanism whereby iASPP control mitotic spindle positioning, we have performed a systematic search for iASPP partners. We have stably expressed a GFP-iASPP construct in SKBr3 cells and fortuitously selected a clone deleted in the C-terminal region, which displayed defective PP1 binding, but strong EB1 binding. Following GFP-trap pulldown, iASPP-associated proteins were identified by mass spectrometry. They comprised a group of proteins known to be associated with the actomyosin cortex^26^ or the plasma membrane (Fig. 4a) including small G proteins and spectrins. The most prominent was myosin-Ic (Myo1c) (Fig. 4b), a member of class I myosin molecular motors, involved in regulating membrane tension, cell adhesion, and actin architecture^27^. Co-immunoprecipitation experiments confirmed that Myo1c associated with C-terminally-truncated, but also full length iASPP (Fig. 4c). Comparably to EB1, Myo1c bound iASPP-ΔAnk-SH3 better than iASPP-WT, but via distinct regions: whereas EB1 clearly relied on the ^350^SRIP^353^ motif, Myo1c binding involved both iASPP1-290 and iASPP382-552 regions (Fig. 4d).

**Figure 4.**
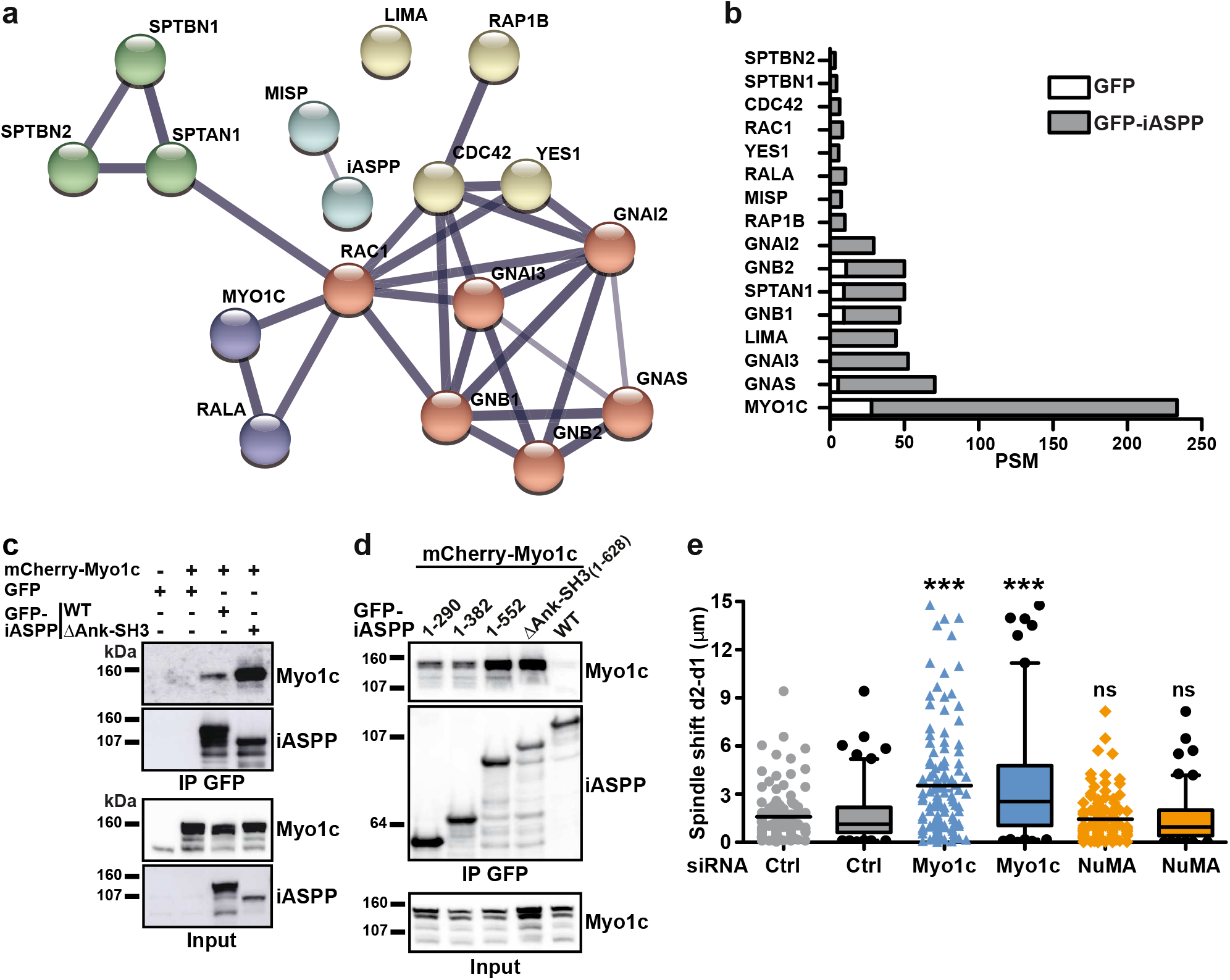
Myosin 1c (Myo1c) is a partner of iASPP, which is also involved in mitotic spindle positioning. (**a-b**) GFP-iASPP was pulled down by GFP-Trap from SKBr3 cells stably expressing a C-terminally truncated form of GFP-iASPP. iASPP-associated proteins were identified by mass spectrometry and those proteins known to localize to the cell cortex or plasma membrane shown as (**a**) a protein-protein interaction network thanks to the STRING database resource (version 11.0) or (**b**) as a bar graph indicating the number of peptide spectrum matches (PSM) identified in the GFP-or GFP-iASPP pulldowns. **(c-d)** Characterization of the regions involved in Myo1c binding. GFP-iASPP constructs and mCherry-Myo1c were expressed in HEK293 cells before immunoprecipitation by GFP-Trap and Western blot analysis. (**e**) Myo1c, but not NuMA, silencing induces abnormal mitotic spindle positioning. HeLa cells were transduced with Ctrl, Myo1c or NuMA targeting siRNA. Mitotic spindle positioning was quantified by comparing the distance between cortex and pole (d1 and d2) on each side of the spindle, as indicated in Fig. 3. Data were pooled from three independent experiments for a total of 120 cells. *** p<0.001; ns: not significant relative to control, using unpaired t-test with Welch’s correction. siCtrl is identical to the one in Fig. 3.

We then tested the involvement of Myo1c in mitotic spindle positioning. In parallel, we evaluated the contribution of the cortically-associated protein NuMA, which is involved in the control of spindle orientation in several cellular and animal models^8,9^. We found that depletion of Myo1c induced a defect in spindle centering comparable to the one induced by iASPP knockdown (Fig. 4e). In contrast, NuMA knockdown had no significant effect on spindle positioning. Thus both iASPP and its partner, Myo1c, contribute to mitotic spindle positioning. The same spindle shift was observed for cells grown on collagen or fibronectin (Fig. 3a, 4e and Supplementary Fig.7a).

Finally, we also evaluated the impact of iASPP and Myo1c on mitotic spindle orientation. We observed that iASPP knockdown, but not Myo1c nor NuMA knockdown, led to abnormal orientation of the mitotic spindle of cells grown on collagen or fibronectin (Supplementary Fig. 7b).

### iASPP and Myo1c contribute to microtubule capture

Because of the cortical localization of the newly identified iASPP partner Myo1c (see Supplementary Fig. 5b), we postulated that the defect in mitotic spindle positioning might be the consequence of abnormal microtubule tethering at the cell cortex. We had previously used live SKBr3 cells, triggered to extend membrane protrusions by the addition of the growth factor heregulin (HRGβ1), as a model to evaluate microtubule capture^2,28–30^. We thus used this model to examine the contribution of iASPP and Myo1c to microtubule capture. We found that in contrast to control SKBr3 cells, where microtubules extended orthogonally to the cell leading edge, towards the cell periphery, iASPP knockdown cells formed microtubules that were parallel to cell leading edge and remained at a distance of the cell cortex (Fig. 5a,b). Ectopic expression of iASPP-WT restored normal microtubule capture (Fig. 5b). In contrast, expression of iASPP-SRNN failed to restore normal microtubule extension, showing that iASPP requires EB1-binding for this function. iASPP-RARA and iASPP-ΔSH3 were equivalent to iASPP-WT, eliminating a potential contribution of PP1 and definitively discarding the involvement of p53. Here again, iASPP-ΔAnk-SH3 was not functional. We similarly evaluated the contribution of Myo1c to microtubule capture. Knockdown of Myo1c also prevented microtubule capture in SKBr3 cells (Fig. 5c). In contrast to re-expression of Myo1c-WT, expression of Myo1c-K389A, mutated in the motor domain, or Myo1c-K892A, mutated in the PH domain (Supplementary Fig. 5b), failed to restore microtubule capture at the leading edge (Fig. 5c), indicating that Myo1c membrane- and actin-binding abilities were required for this function.

**Figure 5.**
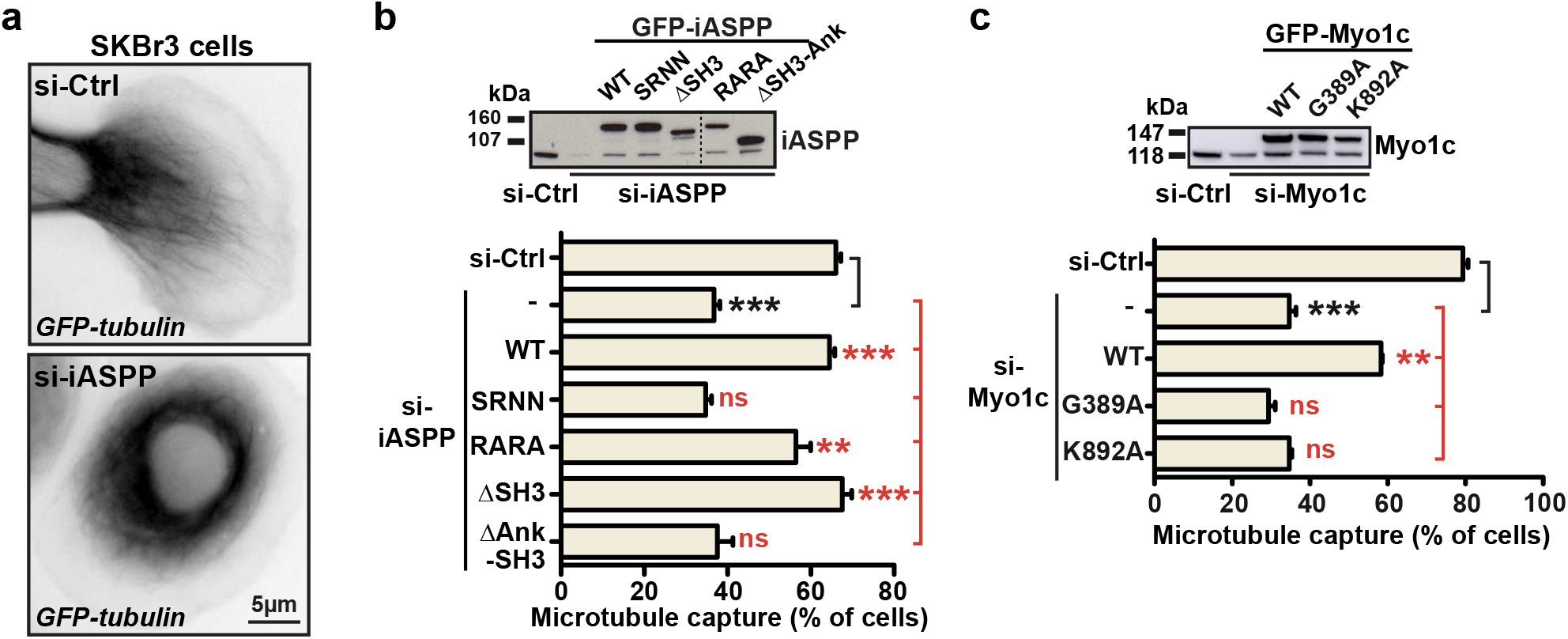
iASPP and Myo1c contributes to microtubule capture at SKBr3 cells leading edge. SKBr3 cells, expressing GFP-tubulin to visualize microtubules, were also transduced with Ctrl, iASPP (**a, b**) or Myo1c (**c**) targeting siRNA and the indicated iASPP (**b**) and Myo1c (**c**) mutated constructs. (**a**) Representative snapshot from video. (**b, c**) The percentage of cells displaying microtubules growing towards the cell cortex was calculated. Bar graphs show mean and SEM from eight (for si-Ctrl, si-iASPP and si-iASPP+WT), five (for all other data points in (**b**)) or three independent experiments (for all data points in (**c**)). Fifty cells were counted per experiment. Western blots above the graphs illustrate the efficiency of the siRNA and the expression levels of the iASPP and Myo1c constructs. *** p<0.001; ** p<0.01; ns: not significant, relative to si-Ctrl (black asterisks) or relative to si-iASPP (red asterisks), using unpaired t-test with Welch’s correction.

Examination of wide-field microscopy images suggested that iASPP also contributes to astral microtubule capture in mitotic HeLa cells. We thus performed a comprehensive 3D-analysis of astral microtubule-ends, following microtubules and EB1-labeling. This allowed us to visualize all astral microtubule-associated EB1 comets and calculate the number of microtubule plus-ends that reached the cell periphery (Fig. 6a,b). Quantification of EB1 comets in control cells revealed a moderate, but significant, asymmetry in the numbers of EB1 comets approaching the cortex on each side of the spindle (Fig. 6b,c). iASPP or Myo1c silencing induced a strong decrease in the number of EB1 comets at the cell cortex (Fig. 6b), strikingly aggravating the asymmetry in the number of astral microtubules that reached the periphery (Fig. 6c). In fact, upon iASPP or Myo1c knockdown, a majority of cells had very few or no comets at one pole (Fig. 6d), while retaining microtubules at the other pole (Supplementary Fig. 8).

**Figure 6.**
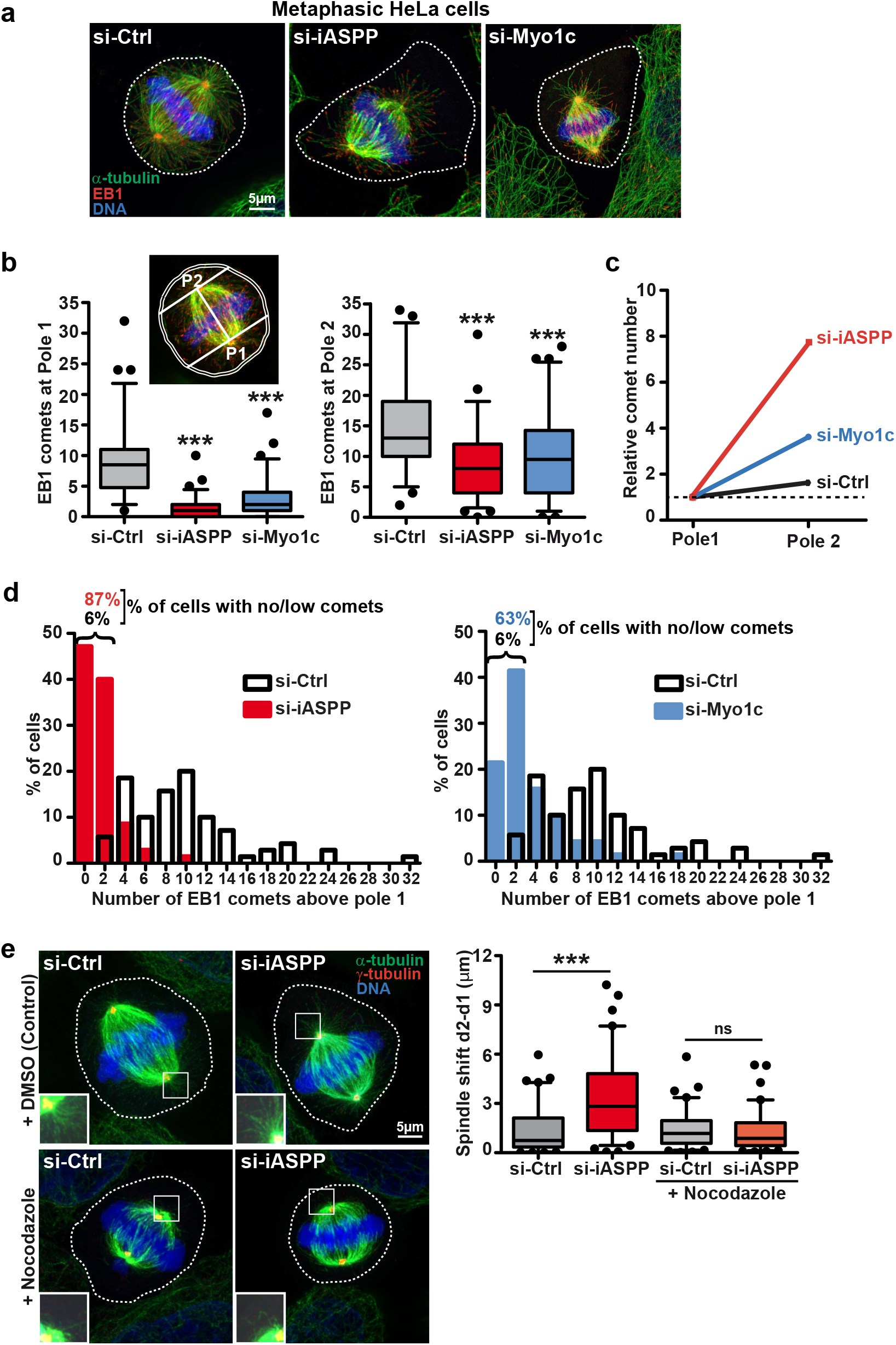
iASPP or Myo1c silencing leads to asymmetrical astral microtubule capture in mitotic HeLa cells. HeLa cells were transduced with Ctrl, iASPP or Myo1c siRNA. (**a**) Microtubules, microtubule plus-ends and chromosomes were visualized by immunofluorescence against α-tubulin, EB1 and DAPI. Forty 0.2 μm sections were collected and a maximum image projection was constructed in order to reveal all astral microtubules and microtubule plus-ends. Representative images are shown. (**b-c**) The number of EB1 comets that entered a zone closer than 2 μm from the cell membrane, at pole 1 (P1, the “less populated” pole) or pole 2 (P2, the “most populated” pole) was counted. Data were pooled from three independent experiments for a total of 70 cells. Data are represented as box-and-whiskers plots with whiskers indicating the 5^th^ and 95th percentile (**b**), as mean comet number normalized to pole 1 (**c**) and as histograms showing the distribution of cells according to comet number at pole 1 (**d**). Mispositioning of the spindle depends on the presence of astral microtubules. HeLa cells, transduced with Ctrl or iASPP siRNAs, were treated with a low concentration of nocodazole in order to induce depolymerization of astral microtubules. Mitotic spindle, centrosomes and chromosomes of metaphasic HeLa cells were visualized by immunofluorescence against α-tubulin and γ-tubulin and DAPI (left). Inserts: zoomed image (1.5x) of the boxed region showing centrosomal area and astral microtubules. Analysis of spindle positioning (right) was performed as in Fig.3. Data were pooled from three independent experiments for a total of 70 cells and presented as box-and-whiskers plots. For all graphs, *** p<0.001, ns: not significant, relative to si-Ctrl using unpaired t-test with Welch’s correction.

Considering the function of astral microtubules in the positioning of the spindle during metaphase, the asymmetry in astral microtubules tethering was likely to be the cause of mitotic spindle off-centering in iASPP knockdown cells. Actually, when we specifically eliminated astral microtubules, at both poles, by pre-treating the cells for a short period of time with low doses of nocodazole, iASPP-silenced cells recovered a properly positioned spindle (Fig. 6e).

### iASPP and Myo1c are required for mitotic cell rounding and proper cortical tension

Upon transition to mitosis, cells disassemble focal adhesions, detach from the substrate and round up because of increased cortical tension, remaining loosely attached through retraction fibers^31^. Unexpectedly, we observed that compared to control cells, iASPP and Myo1c-silenced mitotic cells were not spherical, but were flatter, and more extended in regions close to the substratum (Fig. 7a-c). In fact, iASPP and Myo1c knockdown cells remained juxtaposed to neighboring cells, filling up the space left between interphase cells (Fig. 7a). We measured the maximal length (L) of control and knockdown cells, which was at the level of the spindle axis for control cells, but close to the substratum for knockdown cells (Fig. 7b,c) and calculated the shape ratio (L/W, Fig. 7c) and circularity. The shape ratio was very close to 1 in control cells or cells depleted of NuMA, which were mostly round (Fig. 7d). In contrast, the shape ratio was significantly higher than 1 in both iASPP and Myo1c knockdown cells (Fig. 7d), whereas cell circularity was diminished (Supplementary Fig. 9a). We wondered whether the contribution of iASPP to mitotic cell rounding was dependent on its ability to associate with EB1. We calculated the shape ratio of HeLa cells expressing iASPP constructs defective for EB1 or PP1 binding. We found that cells expressing iASPP-SRNN were significantly misshapen, while cells expressing iASPP-RARA were as spherical as cells expressing iASPP-WT (Fig. 7e). Thus, iASPP, via EB1, and Myo1c contribute to the rounding of cells during the transition to metaphase.

**Figure 7.**
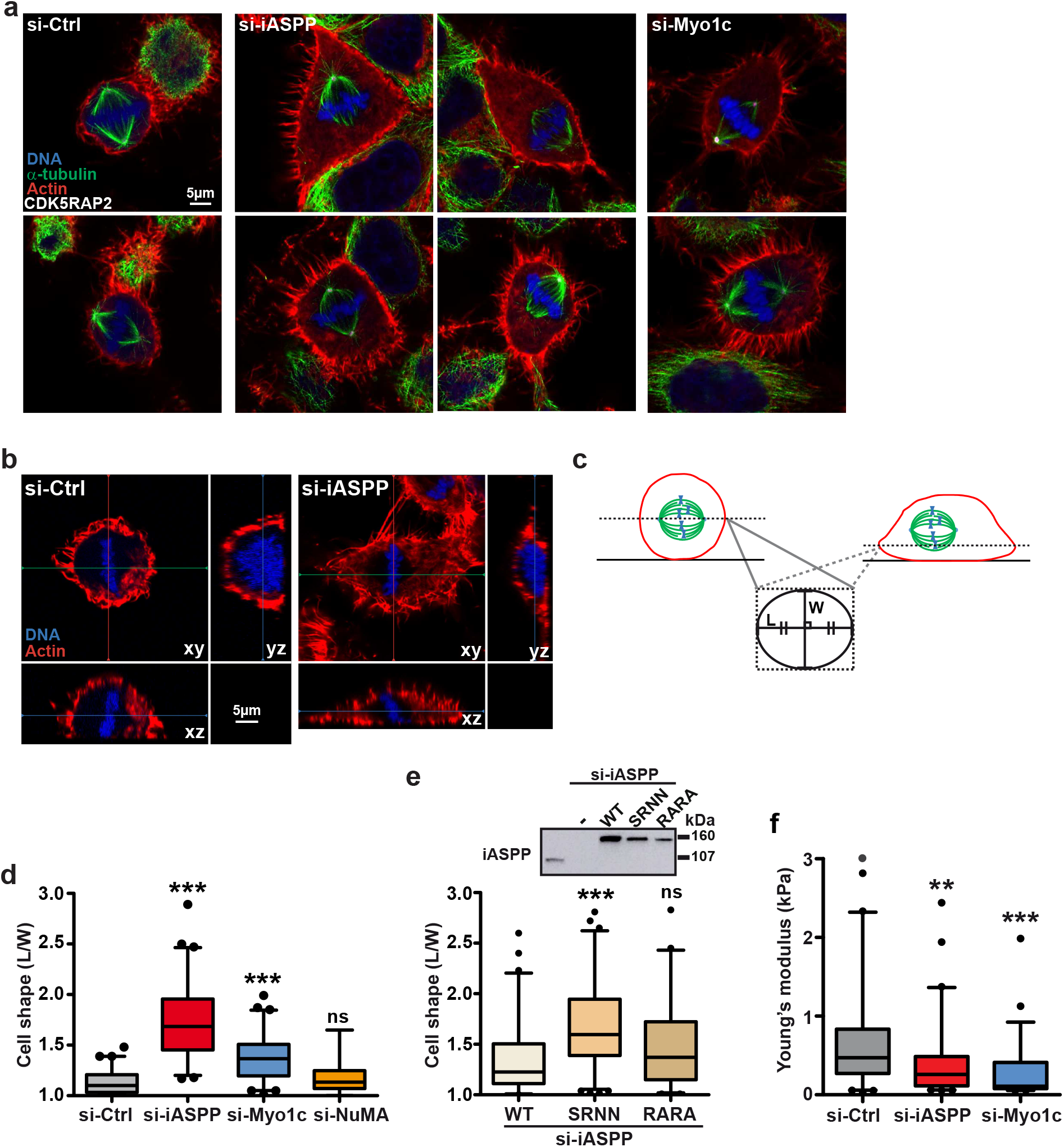
iASPP or Myo1c silencing reduces cortical rigidity and prevents rounding of mitotic cells. (**a-d**) HeLa cells were transduced with Ctrl, iASPP, Myo1c or NuMA siRNA. (**a**) Microtubules, centrosomes, cell cortex, and chromosomes were visualized by immunofluorescence against α-tubulin (**a**) or EB1 (**b**), and CDK5RAP2, TRITC-phalloidin and DAPI staining, respectively. (**a, b**) Representative images are shown. (**b-d**) Changes in metaphase cell shape was quantified by measuring the ratio of the length (L) to the width (W) at the focal plane where the cell is the largest. (**e**) HeLa cells were transduced with iASPP siRNA together with iASPP-WT, -SRNN or -RARA constructs and changes in metaphase cell shape quantified as above. The western blot shows the efficiency of iASPP siRNA and the expression levels of the iASPP constructs. Data were pooled from three independent experiments for a total of 60 cells and presented as box-and-whiskers plots. *** p<0.001, ns: not significant, relative to si-Ctrl in (**d**) or WT in (**e**) using unpaired t-test with Welch’s correction. (f) Stiffness of live mitotic HeLa cells was measured by AFM. Data were pooled from four to five independent experiments for a total of 64, 60 and 55 cells for si-Ctrl, si-iASPP and si-Myo1c, respectively. The grey dot outlier is 3.7 kPa. *** p<0.001; ** p<0.01, relative to si-Ctrl using Kruskal-Wallis with Bonferroni adjustment.

The lacks of cells rounding could be the consequence of inappropriate disassembly of cell-substrate adhesion sites^32^ or defective stiffening of the cortex^12^. Metaphasic iASPP knockdown cells were flat and elongated, but contrary to the neighboring interphase cells, did not display focal adhesions or actin stress fibers (Supplementary Fig. 9b). Thus, iASPP knockdown cells are capable of disassembling focal adhesions when transitioning to mitosis, but yet are unable of rounding. We have therefore investigated the contribution of iASPP and Myo1c to the mechanical properties of HeLa cell cortex in metaphase, using atomic force microcopy (AFM; Fig. S10). We found that when iASPP or Myo1c were silenced, mitotic cell stiffness was greatly decreased, showing a 40 % and 60 % drop in Young’s modulus for iASPP and Myo1 knockdown cells, respectively (Fig. 7f).

## Discussion

We have identified a novel function for iASPP, independent of p53, in the control of astral microtubule capture during metaphase and mitotic spindle positioning. It involves the interaction of iASPP with EB1 at microtubule plus-ends and the control of mitotic cell cortex tension, most likely via Myo1c.

The tumor suppressor p53 is iASPP most investigated partner. In our hands, p53 did not co-precipitate with iASPP. Moreover, iASPP did not show the preferential nuclear localization described originally. This is in accordance with previous data showing that iASPP nuclear localization is restricted to specific cell lines, in particular melanoma^24^ or undifferentiated epithelial cells^19,33^. In addition our results do not support a model in which iASPP is mostly present in the cytoplasm as an anti-parallel homodimer^24^. In fact, our data reveal that iASPP poorly dimerizes, but instead forms a stable complex with PP1. Recent structural studies have identified up to four distinct contact surfaces in the iASPP-PP1 complex that might contribute to complex stabilization. We confirm that iASPP binding to PP1 involves multiple discrete points of interaction and show that, individually, the RARL and SILK motifs, the Ank repeats or the SH3 domain are not sufficient to confer binding, but that altogether they contribute to iASPP-PP1 high affinity interaction. We find that iASPP also binds the microtubule plus-end binding protein EB1, via a typical SxIP motif. In comparison to PP1, we detect only modest amounts of EB1 in iASPP pulldowns; interestingly EB1 interaction with iASPP is enhanced upon loss of PP1 binding.

Our data do not support the formation of intermolecular interactions between iASPP monomers, yet we observe that iASPP N-terminal region can interact with iASPP C-terminal region. The interaction between the N-and the C-termini involves several points of interaction, as was also suggested by a study using peptide arrays^34^, distributed along the disordered region on the one hand, with a particularly important point of interaction located between residues 290 and 382, and within the Ank repeats and the SH3 domain on the other hand. iASPP N-terminal to C-terminal interaction is more easily observed when PP1-binding is prevented. However, preliminary data indicate that even in the absence of PP1 binding, the fraction of iASPP homodimers remains very limited, indicating that the N-terminal to C-terminal interaction is more likely to occur intramolecularly. Future studies should define the structural basis of iASPP intramolecular interactions.

Altogether, we propose that iASPP is able to adopt multiple conformations: a tethered conformation, via intramolecular interactions, and open conformations in association with PP1 or EB1 (Fig. 2h). The equilibrium will depend on relative affinities and local concentration of partners. The affinity of the N-terminal region for the C-terminal region of iASPP was not determined, but the interaction between the overexpressed fragments was considerably competed off by endogenous levels of PP1. The affinity of PP1 for iASPP is in the low nanomolar range^17^, while the affinity of EB1 for SxIP motif-harboring proteins is in the micromolar range^35^ explaining why iASPP-PP1 is the major complex in the cytoplasm. However, EB1 is present at a much higher local concentration at microtubule ends^36^, and in a more favorable conformation for SxIP binding^37^, explaining the recruitment of iASPP to EB1 at this particular location.

It was proposed that phosphorylation of iASPP (at Ser84 and Ser113) prevents iASPP N-terminal to C-terminal interaction^24^, suggesting another level of regulation of iASPP intra/inter-molecular interactions. Global phosphoproteomic studies^38^ and our own mass spectrometry analysis (unpublished observations) confirm that iASPP is highly phosphorylated in the N-terminal region, especially in pseudo-mitotic cells. But we observed that mitotic arrest-induced phosphorylation had no impact on N-terminal to C-terminal interaction, while restraining iASPP-EB1 interaction, calling for careful reexamination of the impact of iASPP phosphorylation. The weakening of iASPP-EB1 interaction upon entry into mitosis could favor the interaction with Myo1c in mitotic cell cortical regions.

Our data show that iASPP or Myo1c knockdown leads to incorrect positioning of the mitotic spindle during metaphase while silencing of NuMA had no impact. Thus, in contrast to most regulators of mitotic spindle positioning/orientation, which act via the control of LGN/NuMA restricted localization^9^, iASPP and Myo1c function via a different mechanism. Several studies describe effectors that potentially impact the proper positioning of the mitotic spindle. These are often connected to EB1 and the control of astral microtubules. For instance, it was shown that knockdown of EB1 or the tumor suppressor APC (adenomatous polyposis coli) induces an increase in mispositioned spindle and a loss of astral microtubules^39^. ASK1 (apoptosis signal-regulating kinase 1) participates in spindle orientation and positioning, by phosphorylating EB1 and stabilizing astral microtubules^40^. Expression of the short isoform of SKAP (small kinetochore-associated protein), mutated in the EB1-binding motif, led to spindles positioned far away from the cell midzone^41^. This was associated with increased lateral growth of astral microtubules along the cell cortex, but only on the side closer to the cortex, conceivably creating force imbalance^41^. In some instances, the effectors were not related to EB1, but rather to the actomyosin cortex: MISP depletion led to movements of the spindle out of the cell center to the cortex^42,43^, which was associated with defective attachment of the astral microtubules to the cortex^43^, even though this was not observed in another study^42^; inactivation of the membrane to cortical actin linker moesin, a member of the ERM family of protein, in Drosophila S2 cells impaired mitotic spindle centering^11^. We observed that iASPP or Myo1c knockdown disturbs astral microtubule capture, weakening attachment to the cortex. It is conceivable that under the constraint of inward directed forces, the weakest cortical attachment (as we have observed that HeLa cells show a small asymmetry in astral microtubule cortical attachment) gives in, while astral microtubules remain associated to the cortex on the other side, providing a force advantage and pulling the spindle that side. Afterwards, few or no astral microtubules will reach the distant edge, as microtubules cannot grow beyond a certain limit^44^, thus maintaining this unbalanced state.

Unexpectedly, iASPP or Myo1c knockdown cells failed to round up upon transition to mitosis. Animal cells undergo dramatic changes in shape and mechanics as they progress through cell division^31^. Mitotic entry is associated with focal adhesions disassembly, osmotic swelling and rearrangement of the cortical actomyosin network, which result in increased cell stiffness and enable cells to adopt a close to spherical shape, even when growing in a crowded environment. The spherical shape provides a suitable environment for the spindle to assemble and limits the space which astral microtubules have to search to connect to the cortex. Rounding also set the stage for the redistribution of masses associated with anaphase and cytokinesis. Mitotic iASPP or Myo1c knockdown cells fail to fully retract, and remain partially extended and juxtaposed to neighboring interphase cells. This could be the consequence of unsuccessful disruption of adhesion sites, as is observed upon overexpression of active Rap1^32^. However, iASPP knockdown cells undergo the expected loss of stress fibers and focal adhesions during the transition to mitosis. Defective rounding is thus most likely to be due to defective cortical tension. Interestingly, upon disruption of the expression or activity of moesin in cultured S2 cells, metaphase cells failed to retract their margins and remained flat^12^, showing strikingly similar images to iASPP knockdown cells. A parallel study showed that moesin knockdown disturbed the mitotic S2 spindle positioning^11^. Importantly, this was associated to strongly reduced cellular stiffness, as measured by AFM^12^. As ERM knockdown HeLa cells do not display the spindle defects or lack of cell rounding^10^, it was suggested that additional cross-linkers play a role in cortical stability in mammalian cells. Class I myosins, including Myo1c, which similarly to ERM proteins connect the plasma membrane to the underlying actin cytoskeleton, are important regulators of membrane tension and cell stiffness in brush border cells, primary mouse fibroblasts and macrophages^45–47^. Our observations demonstrate that Myo1c contributes to cortical stiffness of mitotic HeLa; recent studies indicate that effective stiffness is directly related to cortical tension^48,49^. Unexpectedly, iASPP which does not directly associate with actin or the plasma membrane, had a similar impact on mitotic cell stiffness. Thus, we hypothesize that iASPP functions via the control of Myo1c activity to regulate membrane to cortex attachment which is required for building proper cortex tension; the mechanism whereby iASPP regulates Myo1c activity is an unexplored and exciting field of research. This leads to defective cell rounding upon entry into mitosis, with consequences on astral microtubule attachment to the cell cortex and spindle positioning. It was indeed proposed that a rigid and uniform cortex facilitates physical interactions between the plus-ends of astral microtubules and the actomyosin cortex, enabling the spindle to exert and sustain cortical forces for a proper positioning of the mitotic spindle^12^.

Off-centering of the mitotic cell spindle can lead to unequal daughter cell-size and altered cell cycle progression^50^. Cells have developed mechanisms of correction during anaphase to minimize the impact of this defect^4^. Yet, defective mitotic cell rounding is accompanied by chromosome misalignment, pole splitting, mitotic progression delays or cytokinesis defects, depending on the cellular models^12,32,44^. iASPP knockdown cells showed extended cytokinesis (our preliminary observations and ^51^), but display normal spindle multipolarity and chromosome alignment. However, subtle defects in progression through the cell cycle or chromosomal segregation might only arise after several rounds of divisions. Moreover, the hypertriploid HeLa cells are not the most appropriate model to evaluate the potential impact of iASPP depletion on aneuploidy. The fact that iASPP also affects microtubule capture in migrating cells indicates that, beyond mitosis, iASPP could affect the cell mechanical properties in other physiological situations with possibly much broader consequences.

iASPP was shown to promote tumor development in various *in vivo* cancer models. While this has often been linked to its ability to inhibit p53 pro-apoptotic, this is only relevant to tumors where p53 is not mutated. Considering that novel partners of iASPP have been identified (this study and ^20,51^) and the molecular basis of their interactions determined, tools are now available to evaluate the contribution of iASPP distinct supramolecular complexes to oncogenic progression.

## Methods

### Cell lines

SKBr3 (ATCC/LGC Standards), HEK293F (Thermo Fisher Scientific) and HeLa cells (a kind gift of P. Dubreuil, CRCM, Marseille, France) were cultured in DMEM (Thermo Fisher Scientific) supplemented with 10% fetal bovine serum (Eurobio) at 37°C in a humidified atmosphere with 5% CO2 and were checked regularly for mycoplasma contamination. Plasmid transfection of HEK293F and HeLa cells was performed with Fugene HD (Promega), whereas SKBr3 cells were transfected using Fugene 6 (Promega), following the manufacturer’s instructions. Transfection of siRNA was performed using Lipofectamine RNAiMax (Thermo Fisher) via the reverse transfection method. SKBr3 were co-transfected with cDNA and siRNA by electroporation using Amaxa Nucleofector (kit V, Lonza AG). Cell lysate were collected 48 to 72 hours later. Stable SKBr3 and HeLa cell lines were generated by transfection with the indicated iASPP or Myo1c constructs, selection with 1mg/ml geneticin (Thermo Fisher) and sorting by flow cytometry to select cell populations expressing moderate levels of the transgenes or clones with comparable GFP-tagged construct expression levels. Hela cells expressing GFP-α-tubulin and mCherry-histone H2B constructs were selected successively with geneticin and 0.1 mg/ml hygromycin B.

### Plasmids and siRNA

pEGFP-C1 and pEGFP–α-tubulin were obtained from Clontech. pBabeD-mCherry-histone H2B was a kind gift from C. Lachaud (CRCM, Marseille, France). pDEST-EGFP-C1 and pDEST mycBioID were derived from pEGFP-C1 and pcDNA3.1 mycBioID (a gift from K.J. Roux, Addgene plasmid # 35700) to allow Gateway-based cloning. iASPP cDNA was amplified by PCR from pcDNA3V5iASPP (a gift from X. Lu, Ludwig Institute, Oxford, United Kingdom) and cloned into pDONRZeo. iASPP was subcloned into pDEST-EGFP-C1 in order to obtain GFP-tagged iASPP, pDEST/N-SF-TAP (a gift from C. J. Gloeckner, University of Tübingen, Germany) to obtain strep-Tactin- and FLAG-tagged iASPP (SFII-iASPP) and pDESTmycBioID to obtain BirA-fused iASPP. iASPP point mutations (SRIP to SRNN and RARL to RARA) mutations were produced by Quickchange site-directed mutagenesis (Agilent). Truncated forms of iASPP were generated by PCR amplification and cloned into pDONRZeo, before integration into destination vectors. Plasmid expressing human EB1 (EB1-EGFP JB 131) was a gift from T. Mitchison and J. Tirnauer (Addgene plasmid # 39299). EB1 was cloned into pDEST/N-SF-TAP in order to obtain EB1-SFII and into pcDNA3.1 MCS-BirA(R118G)-HA (a gift from K.J. Roux, Addgene plasmid # 36047) to produce EB1-BirA. Myosin 1c-mCherry construct was a gift from P. Miklavc and M. Frick (Ulm University, Germany), GFP-Myosin1c-WT, -G389A and -K892A constructs were gifts from L. M. Coluccio (University of Boston, MA). All constructs were sequence verified.

The following siRNAs, selected with SVM siRNA Design Tool (Applied Biosystems) software and synthetized by Thermo Fischer Scientific, were used: iASPP-3044 (sense strand: GAAACUUUCCUUAUAAAUATT); Myosin 1c-1480 (sense strand: GGAUAUUUAUGGCUUUGAA); NuMA-7297 (sense strand: GGAUCUUUUCUAAAUGUUA); EB1-1262 (sense strand: UUAAAUACUCUUAAGGCAUTT). An siRNA targeting the β galactosidase gene from E.Coli (LacZ) (sense strand : GCGGCUGCCGGAAUUUACCTT) was used as negative control.

### Antibodies

The following antibodies were used for Western blotting (WB) and immunofluorescence (IF): anti-γ tubulin GTU88 (IF), α tubulin clone DM1A (IF+WB), α tubulin clone YL ½ (IF), β actin clone AC15 (IF), GFP (WB), PPP1R13L HPA041231 (WB), myosin 1c HPA 001768 (IF+WB) from Sigma Aldrich; anti-mCherry (WB) and NuMA [EP3976] (WB) from Abcam; anti-iASPP 18590-1-AP (WB) from Proteintech and PCRP-PPP1R13L-2G4 (WB) from DSHB; anti-EB1 clone 5 (WB) from Cell Signaling Technology; anti-EB1 KT51 (IF), myosin Ic (13) (WB), PP1α (C-19) (WB) from Santa Cruz Biotechnology; anti-NuMA1 clone AD6-1 (IF, WB) from Merck Millipore; anti-CDK5RAP2, A300-554 (WB, IF) from Bethyl Laboratories. GFP-Trap was from Chromotek. Phalloidin-TRITC was from Sigma Aldrich.

### Western Blotting, protein pulldowns and mass spectrometry analysis

Cells were lysed in a Nonidet P-40–based lysis buffer (10mM Tris pH 7.5, 150mM NaCl, 0.5mM EDTA, 0.5% NP-40) supplemented with protease and phosphatase inhibitors (Roche). Cells expressing the GFP-tagged constructs were lysed, and GFP pulldowns were performed using GFP-Trap as described previously^14,29^. Proximity biotinylation was essentially as described ^14,21^. Briefly cells expressing EB1-BirA or BirA-iASPP were treated overnight with 50 μM biotin, were lysed in denaturing buffer (50mM Tris pH 7.4, 500mM NaCl, 0.4 % SDS, 2 % Triton X-100, 5mM EDTA and 1mM DTT) supplemented with protease and phosphatase inhibitors. Biotinylated proteins were isolated by incubating the cell lysates with Avidin-coated beads (ThermoFisher Scientifc) for 1h at 4°C. For Western blotting, samples were run on Novex NuPAGE Bis-Tris 4–12% gels using a 3-(N-morpholino)propanesulfonic acid (MOPS)-based running buffer. Proteins were transferred onto nitrocellulose membranes, incubated with the primary antibodies listed above and secondary antibodies coupled to HRP, and detected by chemiluminescence. Quantification of Western blot was performed with Image J software. For mass spectrometry analysis of GFP-iASPP pulldowns, samples were analyzed by liquid chromatography (LC)–tandem mass spectrometry (MS/MS) in an LTQ-Orbitrap-Velos (Thermo Electron, Bremen, Germany) online with a nanoLC Ultimate 3000 chromatography system (Dionex, Sunnyvale, CA). Protein identification and relative intensity–based label-free quantification was processed using Progenesis LC-MS software, version 4.1 (Nonlinear Dynamics, Newcastle, United Kingdom) as described previously^29^. Only proteins identified with at least two peptides in two out of four independent replicates were selected. Among these, proteins associated with the cell cortex were identified through a literature search.

### Microscopy

HeLa cells, transfected with the indicated siRNA, were grown on glass coverslips coated with 25 μg/mL rat-tail collagen I for 72h. When indicated, nocodazole at a final concentration of 40 nM was added to the culture medium for 1h. For immunofluorescence, cells were fixed either with methanol containing 1 mM EGTA at −20 °C for 5 min, followed by 4% formaldehyde in PBS for 15 min at room temperature; or with 4% formaldehyde and 3% sucrose in PBS for 20 min at 37°C. After permeabilization with 0,2% Triton in PBS for 10 min, immunolabeling was performed with antibodies against target proteins and secondary antibodies labeled with Alexa Fluor 488, 546, 594 or 647 (Jackson ImmunoResearch) and DNA counterstained with DAPI (Sigma Aldrich). Images were acquired on a Zeiss structured light ApoTome microscope equipped with a 63x/1.4 plan Apochromat objective and an Axiocam MRc5 camera using AxioVision software or a Zeiss LSM880 META confocal microscope equipped with a 63x/1.46 plan Apochromat objective and a GaAsP detector using Zen software.

In order to determine spindle position in the *xy* plane, microtubules, centrosome and DNA were labeled as indicated above to identify mitotic cells. Images were acquired with a Zeiss structured light ApoTome microscope. Only metaphase cells with both poles in the same focus plane were quantified, in order to clearly distinguish between effects on spindle positioning and orientation. Cell size, mitotic spindle size, cortex to pole distances (d1 and d2) and cortex to metaphase plate distances (D1 and D2) were measured via Image J software. At total of 150 cells (except when indicated) were measured in three independent experiments. To evaluate spindle orientation along the Z axis, 0.32 μm sections were acquired with a Zeiss confocal LSM880 META, 63x 1.46 plan Apochromat objective and a GaAsP detector, centrosomes position was determined with Zen software and the angle between the line connecting the two centrosomes and the substrate calculated. At least 300 cells were evaluated in three independent experiments.

Astral microtubule plus-ends were identified thanks to EB1 labeling, and quantified on z-stack image projections (40 × 0.2 μm thick sections), acquired with a Zeiss confocal LSM880 META microscope, using a 63x objective and a numeric zoom of 3.5. Astral microtubule EB1 comets that reached an area within 2 μm from the cell periphery, above the poles (as indicated in Fig. 6) were counted, in at least 120 cells in three independent experiments.

To evaluate metaphase cell morphology, cells were labeled with anti-β-actin or TRITC phalloidin, anti-α-tubulin and anti-CDK5RAP antibodies and DAPI, before acquisition of 0,42 μm-thick sections encompassing the entire cell. Length (L) and width (W) of the largest section were determined with Zen software. The largest section was close to the equator in control and NuMA KD cells and two-three sections above the substrate in iASPP and Myo1c KD cells. Circularity was measured in independent experiments: 0.25 μm-thick sections were collected with a ZEISS Axio Observer Z1 microscope equipped with a Yokogawa CSU-X1A head and a plan Apochromat x63/1,46, and cytosolic α-tubulin immunostaining was used to define cell limits using ImageJ particles analysis tool: maximum projection images, blurred with the Gaussian filter (sigma=2 pixels), were used for thresholding, producing a binary image. A 150 pixels^2^ size filter was applied before analysis of cell area, perimeter and circularity [4π(area)/perimeter^2^].

To analyze microtubules in migrating cells^28–30^, SKBr3 cells co-transfected with EGFP-α-tubulin and the indicated siRNA or cDNA were grown on collagen-coated glass coverslips for 48 h and observed 10 min after the addition of 5 nM HRGβ1 (R&D Systems) using the 63X objective (plan Apochromat NA 1.4) of a fluorescence microscope (Zeiss Axiovert 200) driven by MetaMorph 6.3 software. Images were acquired using a CoolSNAP HQ digital camera (Roper) and the percentage of cells showing microtubules extending towards the cell periphery and perpendicular to the cell membrane was calculated in three independent experiments.

### Atomic Force microscopy

EGFP-tubulin/mCherry H2B-Hela cells were transfected with the indicated siRNA, plated on collagen I-coated glass-bottom FluoroDishes (WPI) and synchronized by a single 2mM thymidine treatment. AFM measurements were carried out in serum-free DMEM, 12 to 14 hours after thymidine washout. Details of the setup have been described elsewhere^52,53^. In short, measurements were conducted with an AFM (Nanowizard I, JPK Instruments/Bruker) mounted on an inverted microscope (Zeiss Axiovert 200) with a CellHesion stage and a PetriDish Heater (JPK Instruments/Bruker). The set-up was used in AFM force mode (15 μm head piezo on, stage off) and in closed loop, constant height feedback mode. The sensitivity of the optical lever system was calibrated on the Petri dish glass substrate; the cantilever (Bruker MLCT-bio C) spring constant was determined using the thermal noise method thanks to JPK SPM software routines, at the start and at the end of each experiment. Spring constants were found to be consistently close to the manufacturer’s nominal values (10 pN/nm). Bright field images and/or fluorescence images were acquired with 10X or 40X NA 0.75 lenses, a CoolSnap HQ2 camera (Photometrics) and a four diode Colibri2 (Zeiss) setup with suitable multiband filter sets^52^.

The AFM tip was positioned over the center of mitotic cells (identified by morphology and/or organization of the mitotic spindle). A minimum of 5 force curves were gathered for each cell with a maximal pressing force of 500 pN, a pushing and pulling speed of 2 μm/s and an acquisition frequency of 2048 Hz, making sure that cells were not moving during the indentation measurements with the optical microscope. Experiments were carried at ~ 37°C.

### AFM data processing

Using custom made Python-scripts and JPK-DP data processing software (JPK Instruments/Bruker), force curves were processed. We corrected for baseline offset and tilt, and calculated tip sample separation before applying a fit based on the Hertz model for a square based pyramid^53^. We took care of keeping the fitted indentation to be smaller that minimum manufacturer-reported tip height (< 2μm) and of verifying by eye the goodness of the detection of the contact point for each curve. If the fit appeared of bad quality, the corresponding force curve was rejected from the analysis. We then obtained for each processed curve a Young modulus (in Pa), and calculated the mean Young modulus per cell. The higher the value of Young modulus is, the stiffer the cell is.

### Statistical Analysis

All statistical analyses were performed using GraphPad Prism or R, asbio package, (https://www.r-project.org/). The unpaired two-tailed t-test with Welch correction or Kruskal-Wallis test with Bonferroni adjustment were used to determine if the difference was significant between data groups. Graphs were plotted using Prism, to show the median and/or mean and SEM. P values are indicated on the graphs.

## Data availability

The authors declare that the data supporting the findings of this study are available within the paper and its Supplementary Information files.

## Acknowledgments

We thank M. Rodrigues (CRCM Microscopy and Scientific Imaging Platform) and M. Richaud (CRCM Cytometry Platform) for support. We thank L.M. Coluccio, C. J. Gloeckner, X. Lu, P. Miklavc and M. Frick, T. Mitchison and J. Tirnauer, and K.J. Roux for sharing reagents. This work was supported by Agence Nationale de la Recherche Grant ANR-16-CE11-0008. AM was supported by a Ministère de l’Enseignement Supérieur et de la Recherche Fellowship. The Marseille Proteomics core facility was supported by IBISA (Infrastructures Biologie Santé et Agronomie), Plateforme Technologique Aix-Marseille, Canceropôle PACA, Région Sud Provence-Alpes-Côte d'Azur, Fonds Européen de Développement Régional (FEDER) and Plan Cancer. CRCM Microscopy and Scientific Imaging Platform was supported by FEDER.

## Author contributions

A.M., D.S., M.L.B., S.Q., D.I., S.A., P.H.P. designed and performed experiments and analyzed the data. A.B. and P.V.P. designed the research, analyzed and interpreted the data. AM and AB prepared the figures and A.B. wrote the manuscript, with editing by all the authors.

## Supplementary figure legends

**Figure S1 ǀ** (**a**) **Interaction of iASPP with PP1 requires the RARL motif, the Ank repeats and the SH3 domain.** The indicated GFP-tagged iASPP constructs were co-expressed in HEK293 cells, before immunoprecipitation with GFP-Trap and Western blotting of the GFP constructs and PP1. (**b**) **iASPP phosphorylation does not prevent the interaction between the N-terminal and the C-terminal regions.** HEK293 cells co-expressing SF-tagged N-terminal constructs and GFP-tagged C-terminal of iASPP were treated overnight with nocodazole, before immunoprecipitation with GFP-Trap and Western blotting. The dashed lines indicate a reorganization of the Western blot for the sake of simplicity. (**c**) **iASPP phosphorylation decreases its interaction with EB1.** HeLa cells expressing GFP-iASPP together with EB1-SF or mCherry-Myo1c, were treated overnight with nocodazole, before immunoprecipitation with GFP-Trap and Western blotting of iASPP, EB1, Myo1c and phospho-Histone H3, a marker of mitosis. Note the apparition of a slow migrating band for endogenous iASPP, due to hyperphosphorylation, upon treatment with nocodazole.

**Figure S2 ǀ All iASPP constructs, except GFP-iASPP-SRNN, co-localizes with EB1 at microtubule plus-ends.** Microtubules and EB1 were visualized by immunofluorescence in SKBr3 cells expressing GFP-iASPP, GFP-iASPP-SRNN, GFP-iASPP-RARA, GFP-iASPP-ΔSH3 or GFP-iASPP-ΔAnk-SH3. Inserts: zoomed images (3x) of the boxed region.

**Figure S3 ǀ Efficiency of iASPP, Myo1c, NuMA and EB1 siRNAs.** HeLa cells were transfected with the indicated siRNA, before Western blotting analysis with the corresponding antibodies (**a**) and quantification (**b**). α-tubulin was used as loading control. The bars graph shows mean of three independent experiments and SEM. Signal intensity was normalized to the signal intensity of α-tubulin and Ctrl siRNA was set to 100%.

**Figure S4 ǀ Impact of iASPP silencing on mitotic HeLa cells.**(**a**) The mitotic spindle, centrosomes and chromosomes of Ctrl and iASPP knockdown HeLa cells were immunostained with α-tubulin and γ-tubulin antibody and DAPI. The percentage of metaphasic cells that had a normal (bipolar) spindle, an abnormal (three poles or more) spindle, or chromosomes that failed to align at the metaphasic plate was quantified. Bras show mean with SEM; 80 cells were counted per experiment in three independent experiments. ns: no significant, using unpaired t-test with Welch’s correction. (**b**) **Schematic showing the average control HeLa cell**(black) from 128 cells in three independent experiments; and the average iASPP knockdown HeLa cell (red) from 133 cells in three independent experiments. (**c**) Histograms of the cell distribution of pole-to-cortex distances in control cells (left) and iASPP knockdown cells (right) showing that iASPP knockdown induce a strong d1-d2 asymmetry. (**d-e**) Effect of iASPP silencing on cell diameter (at the poles) and mitotic spindle length. Histograms of cell distribution (**d**) and box-and-whiskers plots (**e**) of cell diameters (left) and spindle lengths (right). *** p<0.001; ** p<0.01, using unpaired t-test with Welch’s correction.

**Figure S5 ǀ Localization of iASPP and Myo1c constructs.** (**a**) **All iASPP constructs, except GFP-iASPP-SRNN, co-localizes with EB1 at microtubule plus-ends in HeLa cells.** EB1 was visualized by immunofluorescence in cells expressing GFP-iASPP, GFP-iASPP-SRNN, GFP-iASPP-RARA, GFP-iASPP-ΔSH3 or GFP-iASPP-ΔAnk-SH3. Inserts: zoomed images (3x) of the boxed region. (**b**) **Localization of Myo1c constructs.** Microtubules and DNA were visualized by immunofluorescence against α-tubulin and DAPI in HeLa cells expressing GFP-Myo1c-WT, GFP-Myo1c-K892A and GFP-Myo1c-K389A. Myo1c-WT and Myo1c-K389A localize to the cell cortex. Myo1c-K892A is mislocalized in the cytoplasm.

**Figure S6 ǀ Schematic showing the average HeLa cell stably expressing iASPP-WT, iASPP-SRNN, iASPP-ΔAnk-SH3 or iASPP-RARA**, obtained from 94, 90, 72 and 102 cells in three independent experiments respectively. The iASPP-SRNN cell is very different from the iASPP-WT cell, but overlaps with the iASPP-ΔAnk-SH3 cell; whereas the iASPP-WT cell largely overlaps with the iASPP-RARA cell.

**Figure S7 ǀ iASPP silencing also affects mitotic spindle orientation.** HeLa cells, grown on fibronectin- or collagen-coated coverslips, were transduced with control (Ctrl), iASPP, Myo1c or NuMA targeting siRNA. (**a**) iASPP silencing induces abnormal mitotic spindle positioning of HeLa cells grown on fibronectin. Mitotic spindle positioning was quantified by comparing the distance between cortex and pole (d1 and d2) on each side of the spindle. Data were collected from three independent experiments and a total of 150 cells. (**b**) Centrosomes, microtubules and chromosomes were visualized by immunofluorescence of CDK5RAP2 and α-tubulin and DAPI. Z-stacks of metaphasic HeLa cells grow on collagen (left) or fibronectin (right) were obtained by confocal microscopy in order to determine coordinates of the spindle poles and calculate α, the spindle angle relative to the substrate (**c**). Data were pooled from three independent experiments and a total of 300 cells. iASPP silencing strongly impacted mitotic spindle orientation while Myo1c and NuMA silencing had very modest or no effect. *** p<0.001; ** p<0.01; * p<0.05; ns: not significant relative to si-Ctrl using unpaired t-test with Welch’s correction.

**Figure S8 ǀ Impact of iASPP and Myo1c on EB1 comet number at pole2.**(**a**) HeLa cells were transduced with Ctrl, iASPP or Myo1c siRNAs. (**a**) Microtubules, microtubule plus-ends and chromosomes were visualized by immunofluorescence against α-tubulin, EB1 and DAPI and the number of EB1 comets that entered a zone closer than 2 μm from the cell membrane, above pole 2 was counted, as indicated in Fig. 6. Data were pooled from three independent experiments for a total of 70 cells and displayed as histograms showing the distribution of cells according to comet number at pole 2. iASPP and Myo1c silencing induced a decrease in comet number, but most cells still had cortical comets at pole 2 (in contrast to pole 1).

**Figure S9 ǀ**(**a**) **iASPP silencing prevents rounding of mitotic cells.** HeLa cells were transduced with Ctrl or iASPP siRNA. Microtubules and chromosomes were visualized by immunofluorescence of α-tubulin and DAPI staining, z-stacks were collected and cell circularity calculated from maximum intensity projection images. Data were pooled from three independent experiments for a total of 186 and 176 cells for si-Ctrl and si-iASPP, respectively and displayed as box-and-whiskers plots. *** p<0.001 relative to si-Ctrl using unpaired t-test with Welch’s correction. (**b**) **Mitotic iASPP knockdown cells normally disassemble focal adhesions and stress fibers.** HeLa cells were transduced with iASPP siRNA. Focal adhesions, actin filaments and chromosomes were visualized by immunofluorescence of paxillin, TRITC-phalloidin and DAPI staining, respectively. Representative images are shown. Two focal planes are presented to visualize focal adhesions (bottom focal plane) and the metaphase plate (middle focal plane). Note the absence of focal adhesions and stress fibers in metaphase cells, compared to neighboring interphase cells.

**Figure S10 ǀ Utilizing AFM forces curves to investigate mitotic cell mechanics.**(**a**) Top view phase contrast (left) and mCherry-H2B fluorescence (right) micrographs showing a cantilever tip positioned near a mitotic cell. Note the typical mitotic cell morphology and chromosome alignment (arrows). (**b**) Typical pressing and pulling force curves. Elastic measurements (Young modulus) were extracted from a fit using a Hertz-like model (green curve).

