## Supplemental Figures for "iASPP contributes to cortex rigidity, astral microtubule capture and mitotic spindle positioning"

Figure S1: Mangon et al

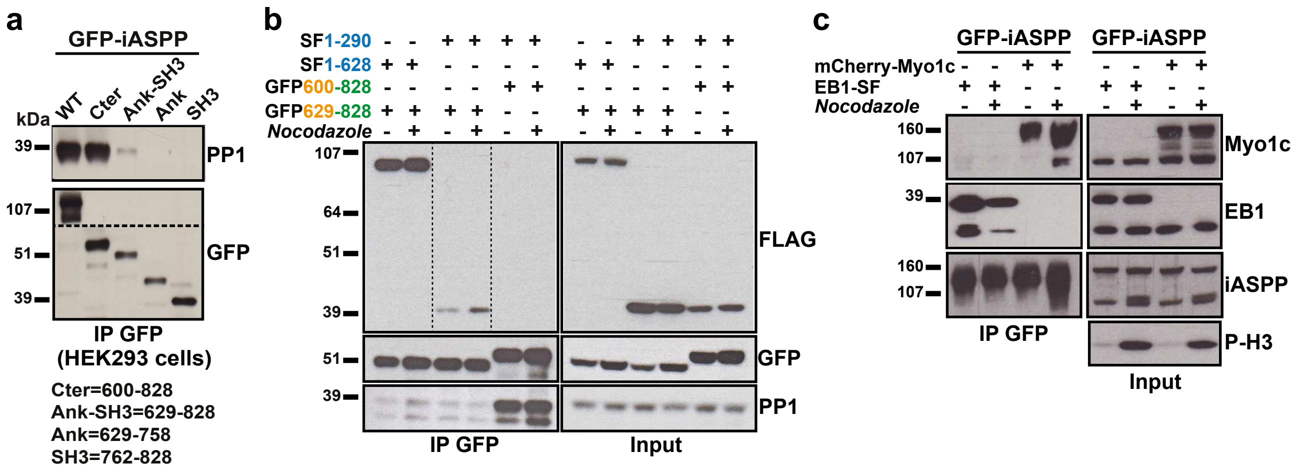


**Figure S1 | (a) Interaction of iASPP with PP1 requires the RARL motif, the Ank repeats and the SH3 domain.** The indicated GFP-tagged iASPP constructs were co-expressed in HEK293 cells, before immunoprecipitation with GFP-Trap and Western blotting of the GFP constructs and PP1. (**b**) **iASPP phosphorylation does not prevent the interaction between the N-terminal and the C-terminal regions.** HEK293 cells co-expressing SF-tagged N-terminal constructs and GFP-tagged C-terminal of iASPP were treated overnight with nocodazole, before immunoprecipitation with GFP-Trap and Western blotting. The dashed lines indicate a reorganization of the Western blot for the sake of simplicity. (**c**) **iASPP phosphorylation decreases its interaction with EB1.** HeLa cells expressing GFP-iASPP together with EB1-SF or mCherry-Myo1c, were treated overnight with nocodazole, before immunoprecipitation with GFP-Trap and Western blotting of iASPP, EB1, Myo1c and phospho-Histone H3, a marker of mitosis. Note the apparition of a slow migrating band for endogenous iASPP, due to hyperphosphorylation, upon treatment with nocodazole.

Figure S2: Mangon et al

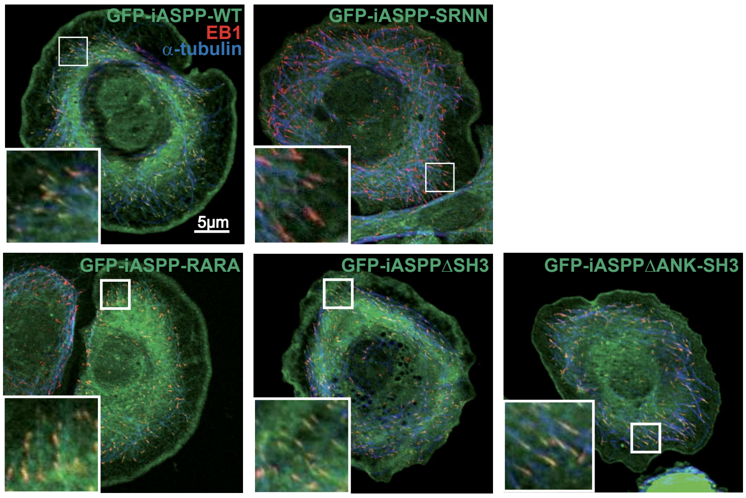


**Figure S2 | All iASPP constructs, except GFP-iASPP-SRNN, co-localizes with EB1 at microtubule plus-ends.** Microtubules and EB1 were visualized by immunofluorescence in SKBr3 cells expressing GFP-iASPP, GFP-iASPP-SRNN, GFP-iASPP-RARA, GFP-iASPP-ΔSH3 or GFP-iASPP-ΔAnk-SH3. Inserts: zoomed images (3x) of the boxed region.

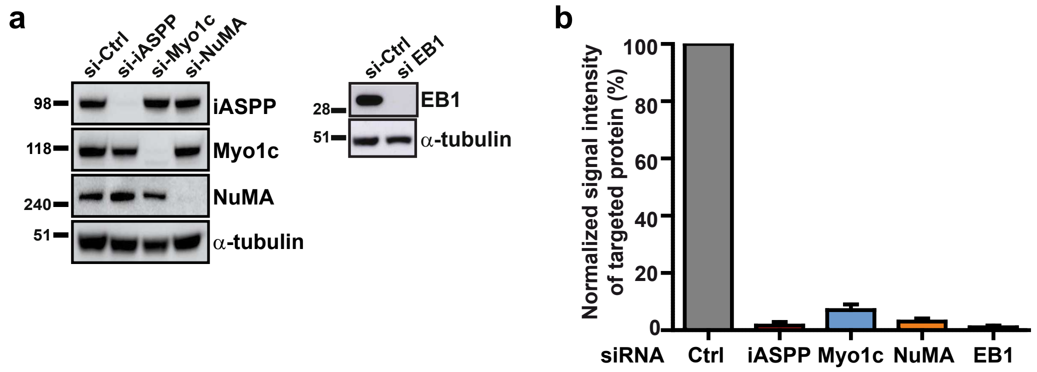


**Figure S3 | Efficiency of iASPP, Myo1c, NuMA and EB1 siRNAs.** HeLa cells were transfected with the indicated siRNA, before Western blotting analysis with the corresponding antibodies (**a**) and quantification (**b**). α-tubulin was used as loading control. The bars graph shows mean of three independent experiments and SEM. Signal intensity was normalized to the signal intensity of α-tubulin and Ctrl siRNA was set to 100%.

Figure S4: Mangon et al

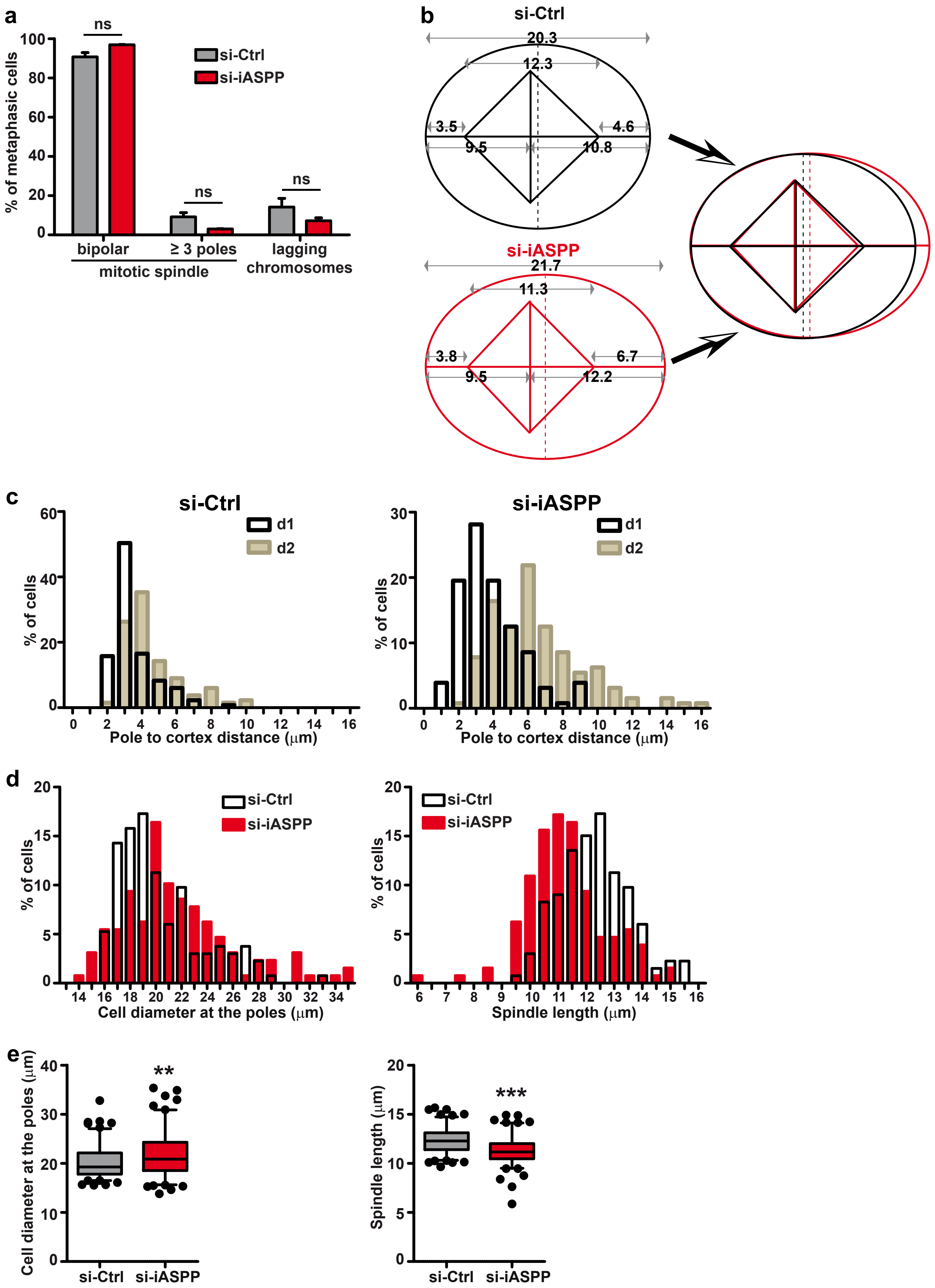

**Figure S4 | Impact of iASPP silencing on mitotic HeLa cells.** (**a**) The mitotic spindle, centrosomes and chromosomes of Ctrl and iASPP knockdown HeLa cells were immunostained with α-tubulin and γ-tubulin antibody and DAPI. The percentage of metaphasic cells that had a normal (bipolar) spindle, an abnormal (three poles or more) spindle, or chromosomes that failed to align at the metaphasic plate was quantified. Bras show mean with SEM; 80 cells were counted per experiment in three independent experiments. ns: no significant, using unpaired t-test with Welch's correction. (**b**) **Schematic showing the average control HeLa cell** (black) from 128 cells in three independent experiments; and the average iASPP knockdown HeLa cell (red) from 133 cells in three independent experiments. (**c**) Histograms of the cell distribution of pole-to-cortex distances in control cells (left) and iASPP knockdown cells (right) showing that iASPP knockdown induce a strong d1-d2 asymmetry. (**d-e**) Effect of iASPP silencing on cell diameter (at the poles) and mitotic spindle length. Histograms of cell distribution (**d**) and box-and-whiskers plots (**e**) of cell diameters (left) and spindle lengths (right). *** $p<0.001$; ** $p<0.01$, using unpaired t-test with Welch's correction.

Figure S5: Mangon et al

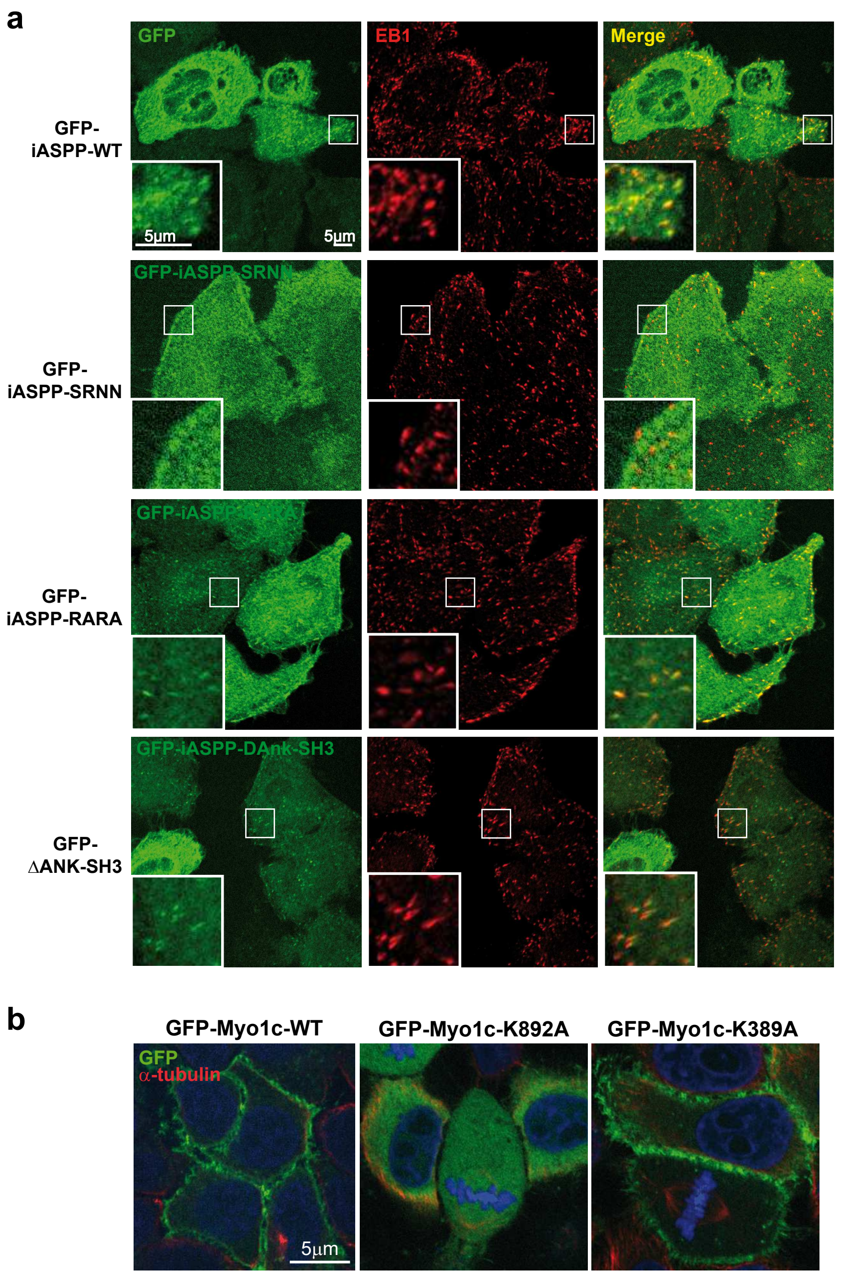


**Figure S5 | Localization of iASPP and Myo1c constructs.** (**a**) **All iASPP constructs, except GFP-iASPP-SRNN, co-localizes with EB1 at microtubule plus-ends in HeLa cells.** EB1 was visualized by immunofluorescence in cells expressing GFP-iASPP, GFP-iASPP-SRNN, GFP-iASPP-RARA, GFP-iASPP-ΔSH3 or GFP-iASPP-ΔAnk-SH3. Inserts: zoomed images (3x) of the boxed region. (**b**) **Localization of Myo1c constructs.** Microtubules and DNA were visualized by immunofluorescence against α-tubulin and DAPI in HeLa cells expressing GFP-Myo1c-WT, GFP-Myo1c-K892A and GFP-Myo1c-K389A. Myo1c-WT and Myo1c-K389A localize to the cell cortex. Myo1c-K892A is mislocalized in the cytoplasm.

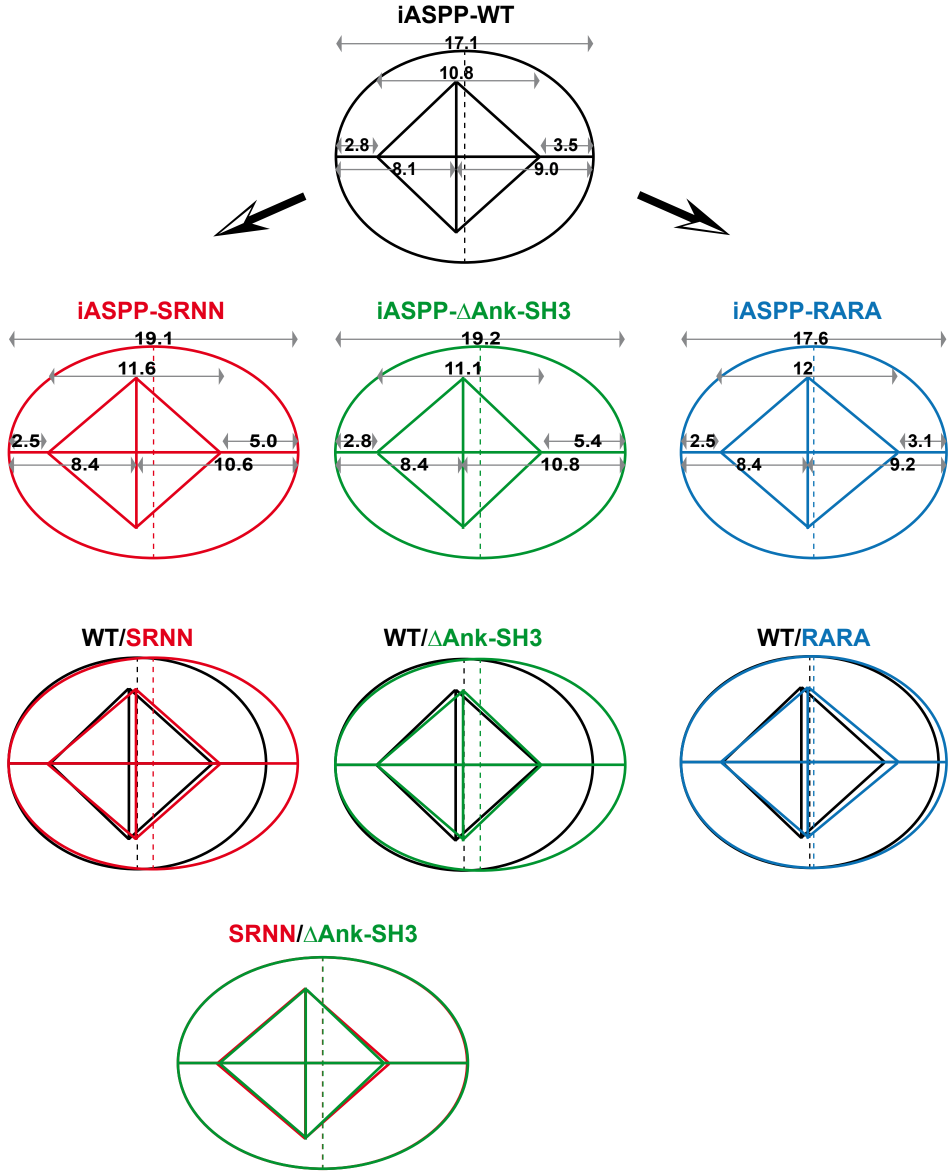


**Figure S6 | Schematic showing the average HeLa cell stably expressing iASPP-WT, iASPP-SRNN, iASPP-ΔAnk-SH3 or iASPP-RARA**, obtained from 94, 90, 72 and 102 cells in three independent experiments respectively. The iASPP-SRNN cell is very different from the iASPP-WT cell, but overlaps with the iASPP-ΔAnk-SH3 cell; whereas the iASPP-WT cell largely overlaps with the iASPP-RARA cell.

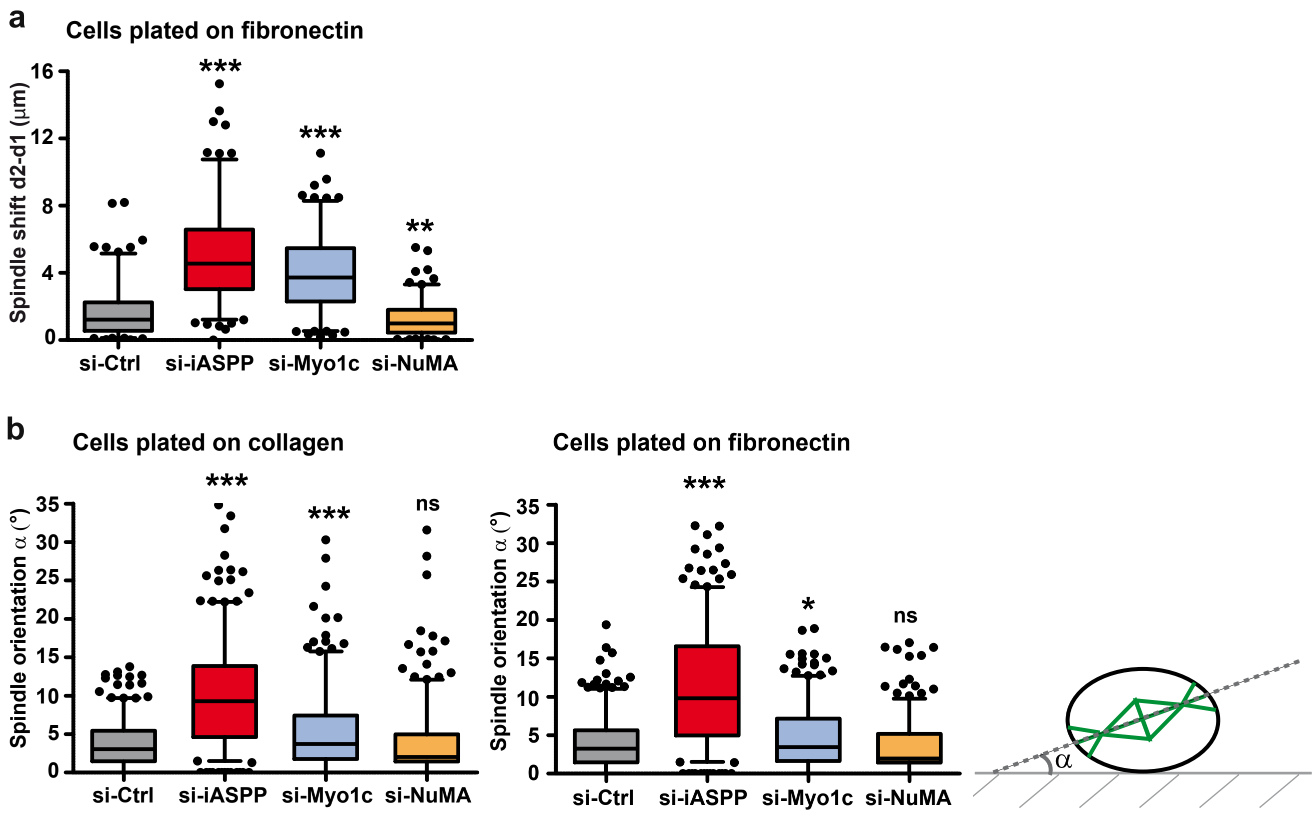


**Figure S7 | iASPP silencing also affects mitotic spindle orientation.** HeLa cells, grown on fibronectin- or collagen-coated coverslips, were transduced with control (Ctrl), iASPP, Myo1c or NuMA targeting siRNA. (**a**) iASPP silencing induces abnormal mitotic spindle positioning of HeLa cells grown on fibronectin. Mitotic spindle positioning was quantified by comparing the distance between cortex and pole (d1 and d2) on each side of the spindle. Data were collected from three independent experiments and a total of 150 cells. (**b**) Centrosomes, microtubules and chromosomes were visualized by immunofluorescence of CDK5RAP2 and α-tubulin and DAPI. Z-stacks of metaphasic HeLa cells grow on collagen (left) or fibronectin (right) were obtained by confocal microscopy in order to determine coordinates of the spindle poles and calculate α, the spindle angle relative to the substrate. Data were pooled from three independent experiments and a total of 300 cells. iASPP silencing strongly impacted mitotic spindle orientation while Myo1c and NuMA silencing had very modest or no effect. *** $p<0.001$; ** $p<0.01$; * $p<0.05$; ns: not significant relative to si-Ctrl using unpaired t-test with Welch's correction.

Figure S8: Mangon et al

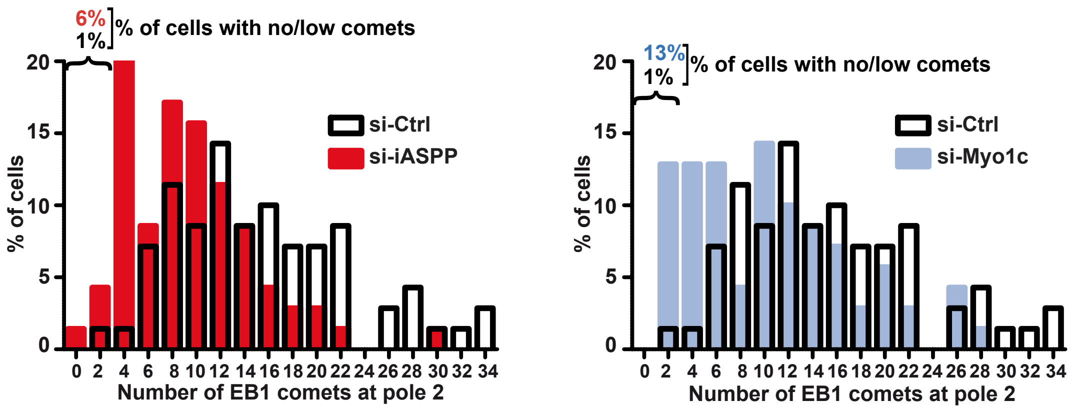


**Figure S8 | Impact of iASPP and Myo1c on EB1 comet number at pole2.** (**a**) HeLa cells were transduced with Ctrl, iASPP or Myo1c siRNAs. (**a**) Microtubules, microtubule plus-ends and chromosomes were visualized by immunofluorescence against α-tubulin, EB1 and DAPI and the number of EB1 comets that entered a zone closer than 2 µm from the cell membrane, above pole 2 was counted, as indicated in Fig. 6. Data were pooled from three independent experiments for a total of 70 cells and displayed as histograms showing the distribution of cells according to comet number at pole 2. iASPP and Myo1c silencing induced a decrease in comet number, but most cells still had cortical comets at pole 2 (in contrast to pole 1).

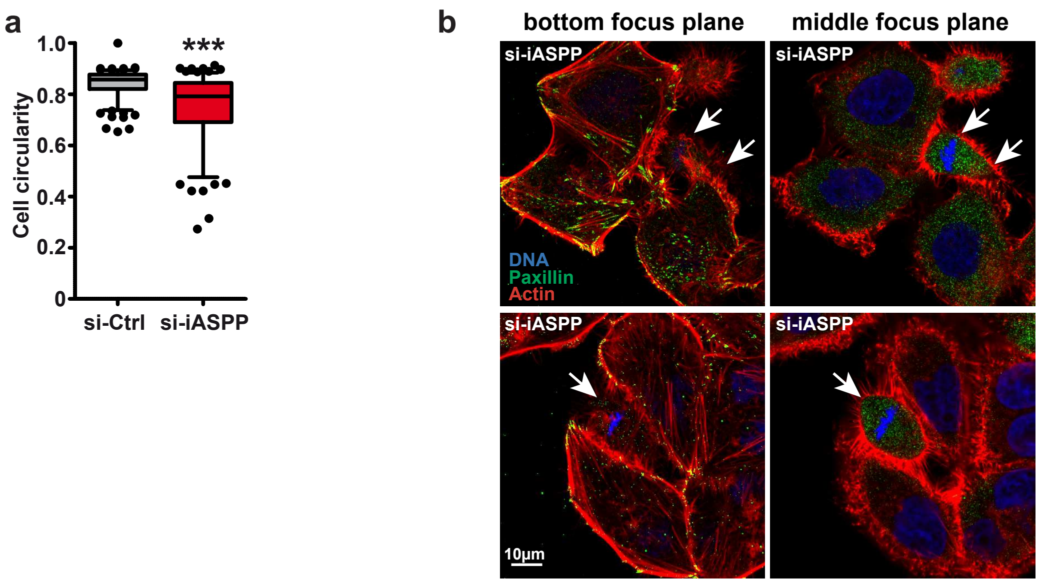


**Figure S9 |** (**a**) **iASPP silencing prevents rounding of mitotic cells.** HeLa cells were transduced with Ctrl or iASPP siRNA. Microtubules and chromosomes were visualized by immunofluorescence of α-tubulin and DAPI staining, z-stacks were collected and cell circularity calculated from maximum intensity projection images. Data were pooled from three independent experiments for a total of 186 and 176 cells for si-Ctrl and si-iASPP, respectively and displayed as box-and-whiskers plots. *** $p<0.001$ relative to si-Ctrl using unpaired t-test with Welch's correction. (**b**) **Mitotic iASPP knockdown cells normally disassemble focal adhesions and stress fibers.** HeLa cells were transduced with iASPP siRNA. Focal adhesions, actin filaments and chromosomes were visualized by immunofluorescence of paxillin, TRITC-phalloidin and DAPI staining, respectively. Representative images are shown. Two focal planes are presented to visualize focal adhesions (bottom focal plane) and the metaphase plate (middle focal plane). Note the absence of focal adhesions and stress fibers in metaphase cells, compared to neighboring interphase cells.

Figure S10: Mangon et al

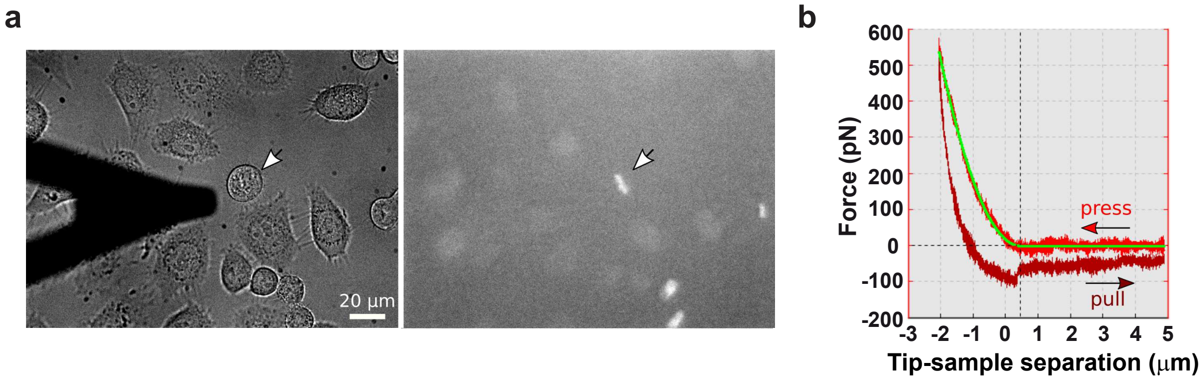


**Figure S10 | Utilizing AFM forces curves to investigate mitotic cell mechanics.** (**a**) Top view phase contrast (left) and mCherry-H2B fluorescence (right) micrographs showing a cantilever tip positioned near a mitotic cell. Note the typical mitotic cell morphology and chromosome alignment (arrows). (**b**) Typical pressing and pulling force curves. Elastic measurements (Young modulus) were extracted from a fit using a Hertz-like model (green curve).
